# Adaptive response to long-term high temperatures during the reproductive development in *Arabidopsis thaliana*

**DOI:** 10.64898/2026.04.24.720607

**Authors:** Juan Francisco Sánchez López, Marie Štefková, Fen Yang, Ales Pecinka, Hélène S. Robert

**Affiliations:** National Centre for Biomolecular Research, Faculty of Science, Masaryk University, 625 00 Brno, Czech Republic; Plant Biotech for Stress Resilience, Genomics and Proteomics for Plant Systems, CEITEC MU - Central European Institute of Technology, Masaryk University, 625 00 Brno, Czech Republic; Institute of Experimental Botany of the Czech Academy of Sciences, Centre of Plant Structural and Functional Genomics, 779 00 Olomouc, Czech Republic

**Author notes:** State Key Laboratory for Crop Stress Resistance and High-Efficiency Production and College of Life Sciences, Northwest A&F University, Yangling, Shaanxi 712100, China (Fen Yang). Juan Francisco Sánchez López. Marie Štefková. Hélène S. Robert. The author responsible for distribution of materials integral to the findings presented in this article is Hélène S. Robert.

**Keywords:** Arabidopsis, Flowering, *hot1*-3, Long-term High temperature, Reproductive development

## Abstract

Increasing global temperatures and the rising frequency of heat waves pose a significant threat to plant reproduction. The reproductive phase is particularly sensitive to heat stress, yet the underlying mechanisms regulating thermotolerance during this stage remain insufficiently understood, despite significant advcances in its understanding during vegetative growth. Heat stress responses are largely controlled by heat shock factors (HSFs) and their downstream targets, including heat shock proteins (HSPs). Among these, HSP101 is essential for acquired thermotolerance and recovery from stress, while HEAT SHOCK BINDING PROTEIN (HSBP) acts as a negative regulator of HSF activity, modulating the heat shock response. Here, we investigated the impact of elevated temperature regimes on the reproductive development of *Arabidopsis thaliana*, with a particular focus on pollen development and fertility. Our results show that heat stress negatively affects pollen development in a dose-dependent manner, leading to reduced reproductive success. We confirmed the critical role of HSP101 in reproductive thermotolerance using the *hot1-3* mutant, deficient in HSP101. Furthermore, we provide evidence that the *hot1-3* mutant is tetraploid. The origin of this event is unknown, but it is tempting to speculate that disruption of heat stress responses and interference with meiotic processes may lead to whole genome duplication. Overall, this study provides new insights into the regulation of plant reproductive development under heat stress and highlights the importance of HSP101 in maintaining fertility. These findings contribute to a better understanding of plant responses to rising temperatures and may inform strategies to enhance crop resilience under climate change.

**Main Conclusion:** Flowering Arabidopsis plants adapt to long-term high temperature by shortening the flowering period and reducing their fertility. The study also demonstrated that the commonly used *hot1-3* mutant is tetraploid.

## Introduction

In recent decades, average global temperatures have increased due to climate change, negatively affecting ecosystems and causing losses amounting to millions of dollars in decreased food and feed production. Climate change not only results in a steady rise in global temperatures but also leads to an increase in the frequency of extreme meteorological events, having a dramatic and adverse impact on crop production. The production of staple crops like rice, wheat, and soybean is estimated to decrease significantly in the upcoming decades due to heat-induced sterility (Khan et al. 2023). High temperatures alter cellular homeostasis in plants, triggering various tolerances. These include the expression of molecular chaperones, detoxifying enzymes such as ascorbate peroxidases, and the production of antioxidants. As sessile organisms, plants cannot escape adverse environmental conditions, leading to complex developmental responses to stresses, including heat. These responses interconnect different hormonal routes, including abscisic acid (Suzuki et al. 2016), cytokinins (Yang et al. 2016; Prerostova et al. 2020), auxin (Ai et al. 2023), ethylene (Jegadeesan et al. 2018), and brassinosteroids (Luo et al. 2022). The hormonal pathways are linked with the expression of molecular chaperones (Shi et al. 1998; Hahn et al. 2011), transcription factors (Nover et al. 2001; Koini et al. 2009; Song et al. 2018; Mizoi et al. 2019), activation of the Unfolded Protein Response (UPR), secondary metabolism and cellular responses (Avin-Wittenberg 2018). However, in severe cases, these adaptive responses may be insufficient to counteract the damage caused by stress conditions. This can result in damaged organs, reduced fertility (Sage et al. 2015; Zhang et al. 2018), and, occasionally, plant death (Vacca et al. 2004; Wahid et al. 2007). Most studies have focused on the vegetative stage of plant development, but the reproductive phase is considered the most sensitive to heat stress (Young et al. 2004; Barnabás et al. 2008; Zinn et al. 2010; Deng et al. 2016; Agho et al. 2024). High temperatures can impair the development of female and male gametophytes, leading to heat-induced sterility (Endo et al. 2009; Zhang et al. 2018; Khaitova et al. 2024) and reducing fertilization of ovules (Snider et al. 2011; Sage et al. 2015), causing smaller fruits after fertilization (Chu & Chang, 2022).

The HEAT SHOCK FACTORs (HSFs) are the primary regulators of the heat shock response (HSR) in plants (Scharf et al. 2012). The most notable HSF is HSFA2, which mediates the HSR (Charng et al. 2007), thermomemory (Friedrich et al. 2021), or other stress responses (Ogawa et al. 2007; Lin et al. 2018b; Zupanska et al. 2019), acting as a super activator (Chan-Schaminet et al. 2009). During the HSR, various HSFs regulate the expression of chaperones (Jacob et al. 2017), with HEAT SHOCK PROTEIN 101 (HSP101) being one of the most studied due to its role in thermotolerance (Queitsch et al. 2000; McLoughlin et al. 2019) and mediating gene expression during the stress recovery phase (Merret et al. 2017). Notably, HSP101 also mediates different aspects of plant reproduction under non-stress conditions (Fu et al. 2002; Qin et al. 2021). The *hot1-3* mutant, a knock-out mutant of the *HSP101* gene (Hong and Vierling 2001), is highly sensitive to heat and often used as a positive control in heat stress experiments (Kim et al. 2012; Wu et al. 2013; Lin et al. 2018a; Tak et al. 2022).

During the heat stress recovery phase, the HEAT SHOCK BINDING PROTEIN (HSBP) acts as a master negative regulator of the HSF transcriptional activity in eukaryotes (Satyal et al. 1998). HSBP binds to the oligomerization domain of HSFs, resulting in the dissociation of active HSF trimers to inactive monomers. The HSF monomers are then coupled to various chaperones (Fu et al. 2005; Rana et al. 2012; Huang et al. 2024). This attenuates the HSF transcriptional activity and HSR (Morimoto 1998). Under non-stress conditions, HSBP is mainly expressed in Arabidopsis siliques (Hsu et al. 2010) and regulates seed development in Arabidopsis, rice, maize and *Brassica rapa* (Fu et al. 2005; Hsu et al. 2010; Rana et al. 2012; Muthusamy et al. 2023). The Arabidopsis *hsbp-2* mutant exhibits a reduced fertilization rate accompanied by about 35% seed abortion (Hsu et al. 2010). HSBP represses HSFs to binding to the promoter of the HEI10 E3 ligase, limiting meiotic crossovers in gametes in normal and high temperatures and, thereby, influencing fertility (Kim et al. 2022). High temperatures alter meiotic crossover formation and induce the formation of unreduced gametes by interfering with cytokinesis during meiotic divisions, producing diploid and aneuploid spores (Storme and Geelen 2020).

This study aimed to provide a comprehensive analysis of the flowering phase of Arabidopsis under different high-temperature scenarios. The results offer new insights into the adaptive response to high temperatures during the plant reproductive phase. We show that high temperatures negatively affect pollen development in a dose-dependent manner and that the *hot1-3* mutant is tetraploid.

## Methods

### Seed Material

We used seeds from Col-0, the GCVC pollen marker line (N67829), *hsbp-2* (salk_046465) (Hsu et al. 2010), *hot1-3* (N16284) (Hong and Vierling 2001), and *hsp101* (N678620/SALK_066374C, N656804/SALK_099583C). All mutant and reporter lines are in the Col-0 accession.

### Growth and Stress Conditions

Arabidopsis seeds were sterilized using chlorine gas for five hours. Seeds were ventilated and plated on Murashige and Skoog (MS) medium. After two days of cold stratification, plates were moved to a cultivation room (Photon Systems Instruments, Czech Republic): 21/18 °C (light/dark) with a 16-hour light/8-hour dark condition (LED illumination) and 50 % humidity. One week after germination, plants were transferred to soil in individual pots. Plants in soil are cultivated in closed cultivation banks in phytotrons with controlled light (combination of white/red/far-red LED illumination) and temperature regimes. Arabidopsis plants remained in this condition (long-day, 21/18) until activation of the inflorescence meristem, corresponding to growth stage 5 (Boyes et al. 2001), before the elongation of the flowering stem started. At this point, plants were moved to different stress conditions or kept in this control condition (CC, 21/18) for the experiments. For Stress Condition 1 (S1, 28/18), the temperature rises from the night temperature (18 °C) to 28 °C in one hour in the first hour of the light period, remains stable for 14 hours, and decreases to 18 °C in the last hour of the light period. For Stress Condition 2 (S2, 28/24), the night temperature is 24 °C and rises to 28 °C in 2 hours at the start of the light period, remains stable for 12 hours, and decreases to 24 °C in the last 2 hours of the light period. For Stress Condition 3 (S3, 34/18), the temperature rises from the night temperature (18 °C) to 34 °C in 5 hours, remains stable for 6 hours, and decreases to 18 °C in 5 hours.

### Phenotyping of Inflorescence

The length of the primary inflorescence of Col-0 plants was measured when the first flower opened in all growth conditions. Measurements were performed in intervals of five days until the end of flowering. Opened and defective flowers were counted daily during the flowering time in each growth condition. Fifteen mature siliques per plant (silique 6 to silique 21, silique 1 being from the first opened flower) from the primary inflorescence were measured to calculate the length of siliques in different conditions. A total of 20 plants were measured for all parameters in three biological replicates.

### Pollen viability and measurements

Pollen grains of 10 plants from all four conditions were stained with Alexander staining (Alexander 1969). Samples were taken 5 and 12 days after the beginning of flowering (DAF, Day After the start of Flowering). One flower was dipped into 15 µL of Alexander staining and vortexed vigorously. Ten microliters of the supernatant were mounted on a slide. Visualization was immediately performed using a ZEISS Axioscope.A1 microscope equipped with DIC optics and an Axiocam 506 CCD camera. The area of 150 pollen grains per plant, ten plants per treatment, was measured using Image J to determine the size of the pollen grains. To quantify pollen viability, at least 200 pollen grains were evaluated. Whole anthers were stained with Alexander staining and kept at 50 °C for 24 hours prior to visualization.

### Pollen ploidy evaluation

Ploidy measurements were performed as described (Yang et al. 2021) with the following modifications. To determine the somatic ploidy levels, 1-2 young leaves from a 10-day-old seedling were chopped with a razor blade in 500 μL Otto I solution (0.1 M citric acid, 0.5 % Tween 20 v/v). The nuclear suspension was filtered through 50 µm nylon mesh and stained with 1 mL of Otto II solution (0.4 M Na_2_HPO_4_·12H_2_O) containing 2 µg DAPI (4’,6-diamidino-2-phenylindole). The ploidy was analyzed on a Partec PAS I flow cytometer with diploid WT plants used as an external standard.

To determine ploidy levels of male gametes, the samples were prepared as described (Borges et al. 2012). The nuclei-enriched pellet was resuspended in 1 mL of LB01 buffer (15 mM TRIS, 2 mM Na_2_EDTA, 0.5 mM spermine, 80 mM KCl, 20 mM NaCl, 15 µM mercaptoethanol, 0.1 % Triton X-100, pH 9) containing 2 µg DAPI. The ploidy was analyzed on FACSAria (Becton Dickinson) flow-sorter with diploid WT male gametes used as an external standard.

To visualize segregating chromosomes, mature pollen samples were prepared as described (Teng et al. 2008).

### Fertilization rate assessment

Pistils from all growth conditions were emasculated and hand-pollinated the following day with pollen of plants from the same genotype grown in 21/18 °C conditions to achieve the maximum pollination rate possible. Siliques were dissected at two, three, and four days after pollination, and developing seeds and unfertilized ovules were counted.

### RNA extraction and cDNA synthesis

Pistils from the primary inflorescence at all growth conditions were emasculated the day before pollination. Forty unpollinated pistils were harvested at midday (at the warmest temperature), frozen on dry ice, and stored at -80 °C. Samples were collected 5 and 12 days after flowering (DAF) began. Samples were ground in liquid nitrogen using ceramic beads and a Retsch Mixer Mill MM 400 (Qiagen). RNA was extracted following the manufacturer’s instructions using the NucleoSpin RNA Plant and Fungi Kit (Macherey-Nagel). After treatment with an RQ1 RNase-Free DNase (Promega), two micrograms of total RNA were used for cDNA synthesis using M-MLV Reverse transcriptase (Promega) following the manufacturer’s instructions.

### RT-qPCR

The FastStart Essential DNA Green Master (Roche) with SYBR Green I as the fluorescent dye was used for qPCR on a LightCycler^®^96 (Roche). Each reaction contained 5 µl FastStart Essential DNA Green Master (Roche), 2.5 µl of four times diluted cDNA sample, 500 nM gene-specific primer combinations, and PCR-grade water (Roche) to a final volume of 10 µL. The selection of the candidate reference genes, TMA7 (At1g15270) and PP2AA3 (At1g13320), for normalization was made using qBASE+ (BioGazelle) and a combination of the Normfinder algorithm in R and BestKeeper Excel sheet (Vandesompele et al. 2002; Andersen et al. 2004; Taylor et al. 2019). The expression of the reference genes was not affected by the growth conditions. The primers targeting the tested transcripts are listed in Supplementary Table S3. Primers for target genes were validated by testing their efficiency and specificity. Three biological replicates were used for the experiments, each with three technical replicates. The PCR reactions consisted of a preincubation step at 95 °C for 10 minutes, followed by 40 cycles of 95 °C for 10 s, 55 °C for 10 s, and 72 °C for 14 s. Products were checked on a 3% agarose gel. The calculation of log2-fold changes was made using the 2^DCq method according to the guidelines in Taylor et al. (2019). Statistical analysis was performed using a two-way ANOVA and post-hoc Tukey test in R Studio. A log2-fold change of 0.8 or higher was considered significant.

### Statistical analysis

To compare flowering times in the different growth conditions, a negative binomial regression test was performed, followed by ANOVA and Tukey test to make multiple pairwise comparisons. A Kaplan-Meier Survival analysis and Cox proportional hazards regression model were used to study the flowering speed, the end of flowering, and the presence of sterile flowers. One-way or two-way ANOVA and Tukey HSD tests were performed to compare the length of siliques and pollen size. To study the viability of pollen grains, number of flowers, number of ovules, number of developed seeds, non-fertilized ovules, and aborted seeds, Kruskal-Wallis, Mann–Whitney U and Dunn tests were performed, followed by a non-parametric Spearman rank correlation to establish a correlation between the number of ovules and developed seeds.

### Accession Numbers

Arabidopsis AGI locus identifiers: APX2 (At3g09640), HSBP (At4g15802), HSFA1D (At1g32330), HSFA2 (At2g26150), HSFA4C (At5g45710), HSP70 (At3g12580), HSP101/HOT1 (At1g74310), PP2AA3 (At1g13320), TAA1 (At1g70560), TMA7 (At1g15270),

## Results

### Impact of long-term high temperatures on flowering rate and flower production

Various phenotypic changes in the flowering plants were notable when we analyze the effects of long-term high temperatures (Supplementary Figure S1). In all tested growth conditions, flowering commenced six to nine days after transferring the plants to the different stress and control conditions. The duration of flowering significantly shortened under high-temperature conditions, decreasing from approximately twenty-one days in the control condition to a range of twelve to fifteen days in all stress conditions (Fig. 1a). This reduction in flowering time affected the growth of the primary inflorescence branch, which we monitored by measuring its length at different time points (Fig. 1b). In the Control Condition (hereafter, referred to as CC), plants uniformly elongated until the end of flowering. Under Stress Condition 1 (28/18 °C, hereafter, referred to as S1) and Stress Condition 2 (28/24 °C, hereafter, referred to as S2), growth accelerated until ten days after the beginning of flowering (10 DAF), then slowed from 10 DAF to the end of flowering. Under Stress Condition 3 (34/18 °C, hereafter, referred to as S3), there was a significant reduction in the inflorescence growth rate compared to the control condition, indicating compromised inflorescence meristem activity.

**Fig. 1.**
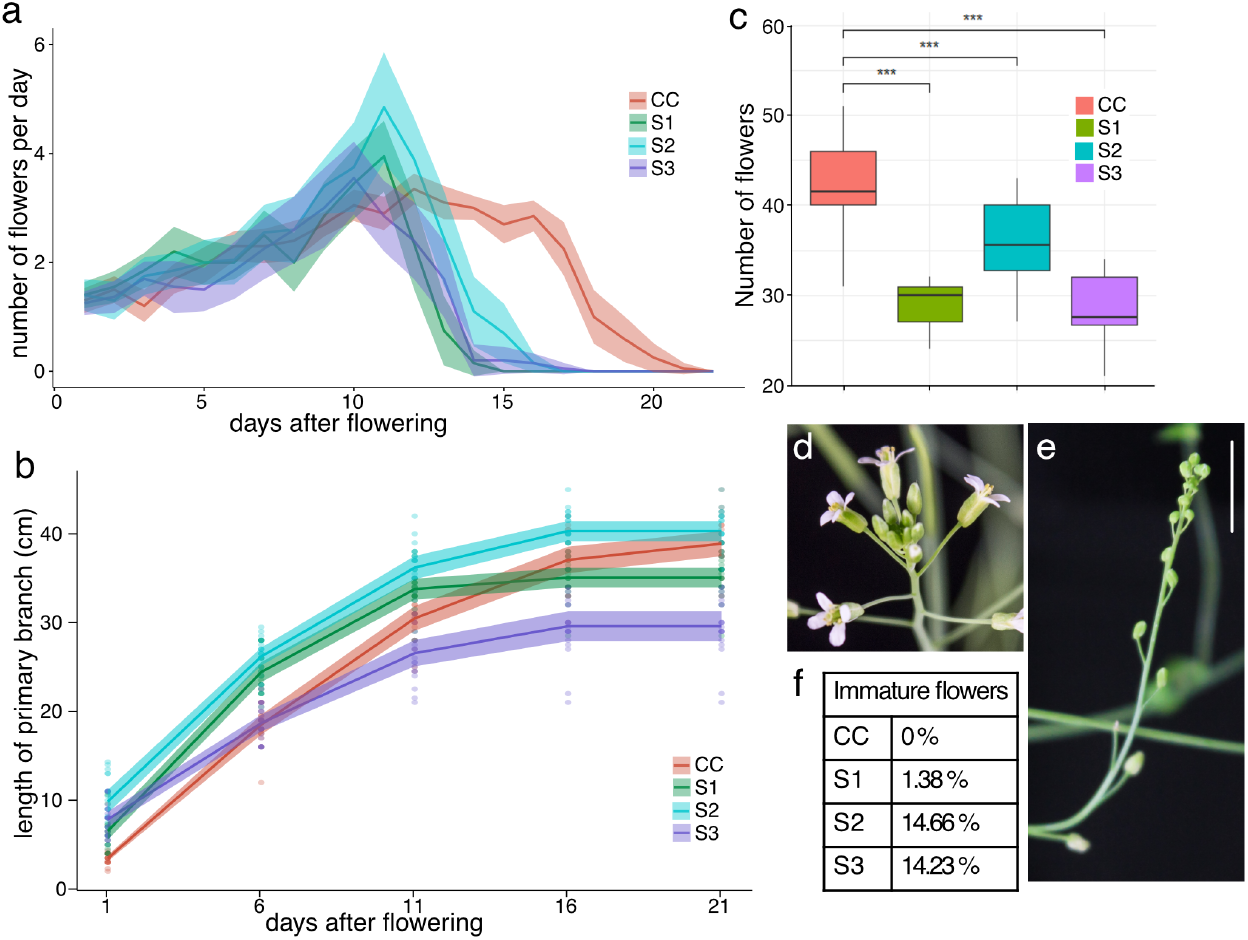
Effect of high temperatures on flower production. (a) Daily production of flowers in each growth condition on the primary inflorescence branch. Statistical analysis of the flowering rate is presented in Supplementary Table S1. (b) Length of the primary branch in cm in each growth condition. The 95% CI and mean are shown in (a) and (b). (c) Total number of flowers produced in the primary branch. Significant difference with p-value of *p* < 0.001 is indicated by ***. (d, e) Representative pictures of normal flowers (d) and immature flowers (e). The scale bar represents 1 cm. (f) Percentage of immature flowers produced per plant (n = 20). Statistical analysis using a Kaplan-Meier survival estimator and Cos proportional hazard regression is presented in Supplementary Fig. S2. CC: Control Condition (21/18 °C); S1: Stress Condition 1 (28/18 °C); S2: Stress Condition 2 (28/24 °C); S3: Stress Condition 3 (34/18 °C).

Reduced flowering duration led to the production of fewer flowers (Fig. 1c). Under CC, plants produced two to four flowers daily on the primary inflorescence branch. In contrast, under stress conditions, plants produced up to eight flowers daily, with a high variability day to day (Fig. 1a). High temperatures also induced the production of flower buds that stopped their development in a dose-dependent manner, up to 14 % at Stress Conditions S2 and S3 (Fig. 1, d-f). This indicates that high temperatures not only reduce the primary inflorescence branch growth rate but also reduce flower production, with a significant proportion of flowers that will not reach maturation under strong stress conditions. Inflorescence development and flower production were most affected in S2 and S3. Our statistical analysis, utilizing an exponential growth model (Supplementary Figure S2), revealed a strong correlation between inflorescence growth and flowering duration across different conditions, suggesting that high temperatures directly impact the flower meristem activity.

### Pollen grains are damaged by high-temperature stress

To better understand how high temperatures influence seed production, we investigated the effects on pollen grains in plants grown under various high-temperature conditions. Pollen is highly sensitive to heat (Rieu et al. 2017; Raja et al. 2019). Using Alexander staining on stamens, we observed that, in CC, approximately 0.1 % (n = 2012) of pollen grains were aborted, while in stress conditions, the percentage of aborted pollen grains in 5 DAF flowers varies from 3 % (n = 2308) in S1, 4.6 % (n = 2364) in S2, to 7.3 % (n = 2021) in S3. Interestingly, in 12 DAF flowers, pollen grains appeared to adapt to the stress conditions with reduced abortion rates of 1.2 % (n = 2270, S1), 3.5 % (n = 2131, S2), and 3.8% (n = 2283, S3) (Table 1, Fig. 2a). This slight decrease in pollen viability at high temperatures compared to control temperature is significant at these time points.

**Table 1.** Pollen viability assessed by Alexander’s staining (n=10 plants) DAF, Days After Flowering; CC, Control Conditions; S1, Stress conditions 1; S2, Stress conditions 2; S3, Stress conditions 3. Statistical analysis was performed by Kruskal-Wallis and Dunn multiple comparison tests with *p*-values adjusted by the Holm method. Non-significantly different samples were grouped by letter. The first letter for comparing the effect of time in each growth condition; and the second letter for comparing the effect of growth temperature at each time point.

| DAF | Condition | aborted pollen<br>(n) | % of aborted<br>pollen | statistical<br>groups |
| --- | --- | --- | --- | --- |
| 5 | CC | 2 (2012) | 0.10 | a, b |
| 12 | CC | 3 (2174) | 0.14 | a, a |
| 5 | S1 | 68 (2308) | 2.95 | a, a |
| 12 | S1 | 28 (2270) | 1.23 | b, a |
| 5 | S2 | 110 (2364) | 4.65 | a a |
| 12 | S2 | 76 (2131) | 3.57 | a, b |
| 5 | S3 | 147 (2021) | 7.27 | a, a |
| 12 | S3 | 88 (2283) | 3.85 | b, c |

**Figure 2.**
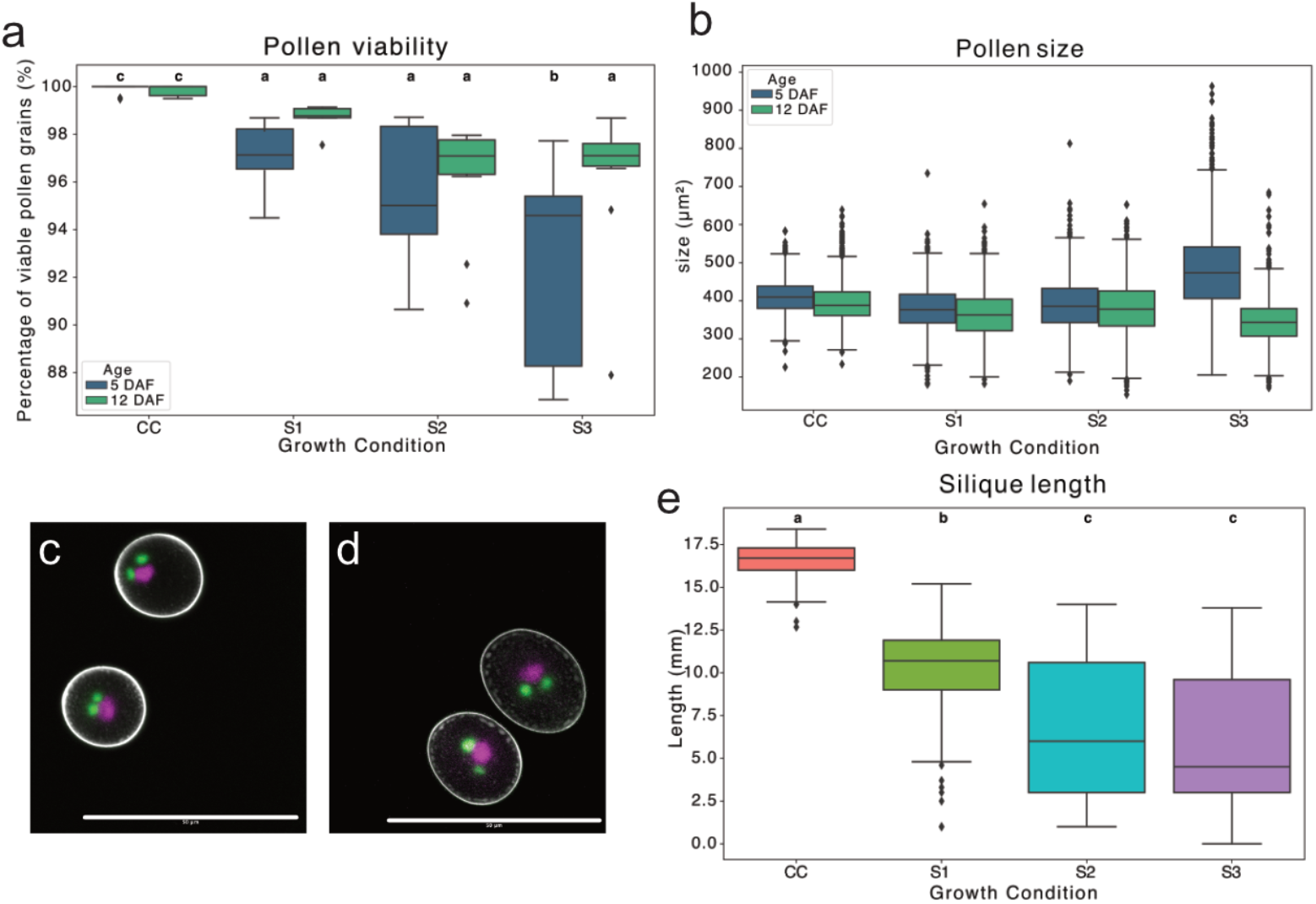
Effect of long-term high temperatures on pollen development. (**a**) Pollen viability assessed by counting the number of viable pollen grains from 10 plants grown under the four growth conditions (CC, control; S1, Stress 1; S2, Stress 2; S3, Stress 3) visualized by Alexander’s stain at 5 and 12 Days After Flowering (DAF) (n=1874 - 2254). Statistical analysis was performed by Kruskal-Wallis, Mann-Whitney U test and Dunn tests. Data are presented as boxplot with median value, interquartile range as box and minimum and maximum values as whiskers. Groups with the same letter are not significant. (**b**) Pollen size in µm^2^ from 150 grains from 10 plants grown under the four growth conditions. Statistical analysis was performed by two-way ANOVA followed by Tukey HSD tests. (**c**) Pollen grains from a SCVC marker plant grown at CC. The sperm cell nuclei are marked in green and the vegetative cell nucleus is marked in magenta. Scale bar is 50 µm. (**d**) Pollen grains from a SCVC marker plant grown at S3. The sperm cell nuclei are marked green and the vegetative cell nucleus is marked in magenta. Scale bar is 50 µm. (**e**) Silique length in mm from self-pollinated flowers (10 siliques) from 15 plants grown under the four growth conditions. Statistical analysis was performed by one-way ANOVA followed by Tukey HSD tests, showing significant differences between groups with *p* < 0.0001. Groups with the same letter are not significant. CC: Control Condition (21/18 °C); S1: Stress Condition 1 (28/18 °C); S2: Stress Condition 2 (28/24 °C); S3: Stress Condition 3 (34/18 °C).

Remarkably, the size of pollen grains also varied between stress conditions. In CC, S1 and S2, pollen grain size was similar, averaging around 400 µm^2^. Notably, a higher number of small pollen grains (around 200 µm^2^) was only observed in stress conditions. Under S3, pollen grains in 5 DAF flowers were significantly larger, sometimes twice the size of those from the CC (Fig. 2b). These large pollen grains were not observed in 12 DAF flowers. Using the fluorescent GCVC reporter line marking distinctly the sperm cell nuclei and vegetative cell nucleus, we found no apparent changes in the number of nuclei in larger pollen grains despite their size (Fig. 2, c and d).

### The fertilization rate is reduced at high temperatures

Self-pollination revealed a dose-dependent reduction in fertilization rates. The fertilization rates significantly dropped from 97.43 % in CC to 79.04 %, 51.31 %, and 44.62 % under S1, S2, and S3, respectively (Table 2). Additionally, seed abortion rates increased to 0.82 % and 1.82 % at S2 and S3, respectively. The correlation between the number of ovules and developed seeds (number of ovules versus number of developed seeds) weakens as temperature rises (Spearman rank correlation rho 0.95 for CC, 0.70 for S1, -0.04 for S2, -0.07 for S3), with S2 and S3 showing the least significant effects, suggesting a negative effect of the long-term high temperature on fertilization rate.

**Table 2.** Fertilization rate. Developed seeds, aborted seeds, and unfertilized ovules were counted in 25 siliques per growth condition. CC, Control Conditions; S1, Stress conditions 1; S2, Stress conditions 2; S3, Stress conditions 3. Statistical analysis was performed by Negative Binomial regression, except for seed abortion (Kruskal-Wallis and Dunn multiple comparison tests with *p*-values adjusted by the Holm method). Aborted seeds are also counted as fertilized ovules. Developed seeds are fertilized ovules that form seeds and do not abort.

| Condition |  | C | S1 | S2 | S3 |
| --- | --- | --- | --- | --- | --- |
| silique | n | 25 | 25 | 25 | 25 |
| fertilized ovules | n | 1741 | 1139 | 627 | 564 |
|  | % | 97.43 | 79.04 | 51.31 | 44.62 |
|  | *p*-values (compared to CC) |  | 0.0365 | <0.0001 | <0.0001 |
| unfertilized ovules | n | 46 | 302 | 595 | 700 |
|  | % | 2.57 | 20.96 | 48.69 | 55.38 |
|  | *p*-values (compared to CC) |  | <0.0001 | <0.0001 | <0.0001 |
| aborted seeds | n | 0 | 0 | 10 | 23 |
|  | % | 0.00 | 0.00 | 0.82 | 1.82 |
|  | *p*-values (compared to CC) |  | 1 | 5.72E-02 | 2.30E-06 |
| total n |  | 1787 | 1441 | 1222 | 1264 |
| n ovules/silique |  | 71.48 | 57.64 | 48.88 | 50.56 |
| *p*-values (compared to CC) |  |  | <0.0001 | <0.0001 | <0.0001 |
| n dev. seeds/silique |  | 69.64 | 45.56 | 24.68 | 21.64 |
| *p*-values (compared to CC) |  |  | 0.0365 | <0.0001 | <0.0001 |

The reduced fertilization rate and increased seed abortion led to significantly fewer developed seeds per silique (Table 2). Under CC, plants averaged 69.64 developed seeds per silique, whereas this amount was reduced to 45.56, 24.68, and 21.64 under S1, S2, and S3, respectively (Table 2). This reduction in seed number also significantly affected the length of the siliques measured at flower stage 17, from 16 mm in the CC to 9.5, 7.9, and 7.4 mm in S1, S2, and S3, respectively (Fig. 2e).

### The HSR is active in temperature-stressed pistils

Heat shock and oxidative responses are crucial in managing temperature stress. To understand the role of the HSR components in response to high temperatures during fertilization, we collected unpollinated pistils at 5 and 12 DAF and analyzed the expression of the various heat-responsive genes across the different growth conditions. We focused on the expression of *HSF* genes known to be expressed in reproductive tissues (*HSFA1d, HSFA2, HSFA4c*), genes related to the HSR (*HSP70, HSP101, APX2, HSBP*), and auxin biosynthesis (*TAA1*) (Fig. 3). The duration of the growth at stress conditions had no impact on the expression of the tested genes, excepted for *HSBP* (Fig. 3d) for which a prolonged warmness led to a significantly more pronounced upregulation. The *TAA1* expression was significantly downregulated in pistils from plants grown under all stress conditions compared to plants grown in CC (Fig. 3h). *HSFA2, HSP70, HSP101*, and *APX2* were the most significantly overexpressed genes in pistils from plants grown at S3 compared to plants grown in CC and S1 and S2 (at 5 DAF, up to 6.3-, 7.0-, 5.1-, and 8.8-log_2_-fold change, respectively, Fig. 3, b, e-g). To a lesser extent, *HSFA1d* and *HSFA4c* genes were also significantly upregulated in pistils from plants grown under S2 and S3 compared to plants grown in CC (at 5 DAF, 1- and 1.1-log_2_-fold change, respectively, Fig. 3, a and c). *HSFA1d, HSFA4c* and *HSBP* were only mildly affected by the growth conditions. It is important to note that *HSF* transcripts can undergo post-transcriptional regulation, affecting their activity and stability in the cell (Cohen-Peer et al. 2010; Evrard et al. 2013; Carranco et al. 2017; Andrási et al. 2019). Therefore, even small upregulations in these transcription factors can significantly impact cellular responses, as they may already be present in an inactivated state. In addition, this gene expression analysis indicated that S3 appears to be the most stressful and that gene expression patterns under S1 and S2 were rather similar. Therefore, we decided to continue our analysis only with plants grown at S1 and S3.

**Figure 3.**
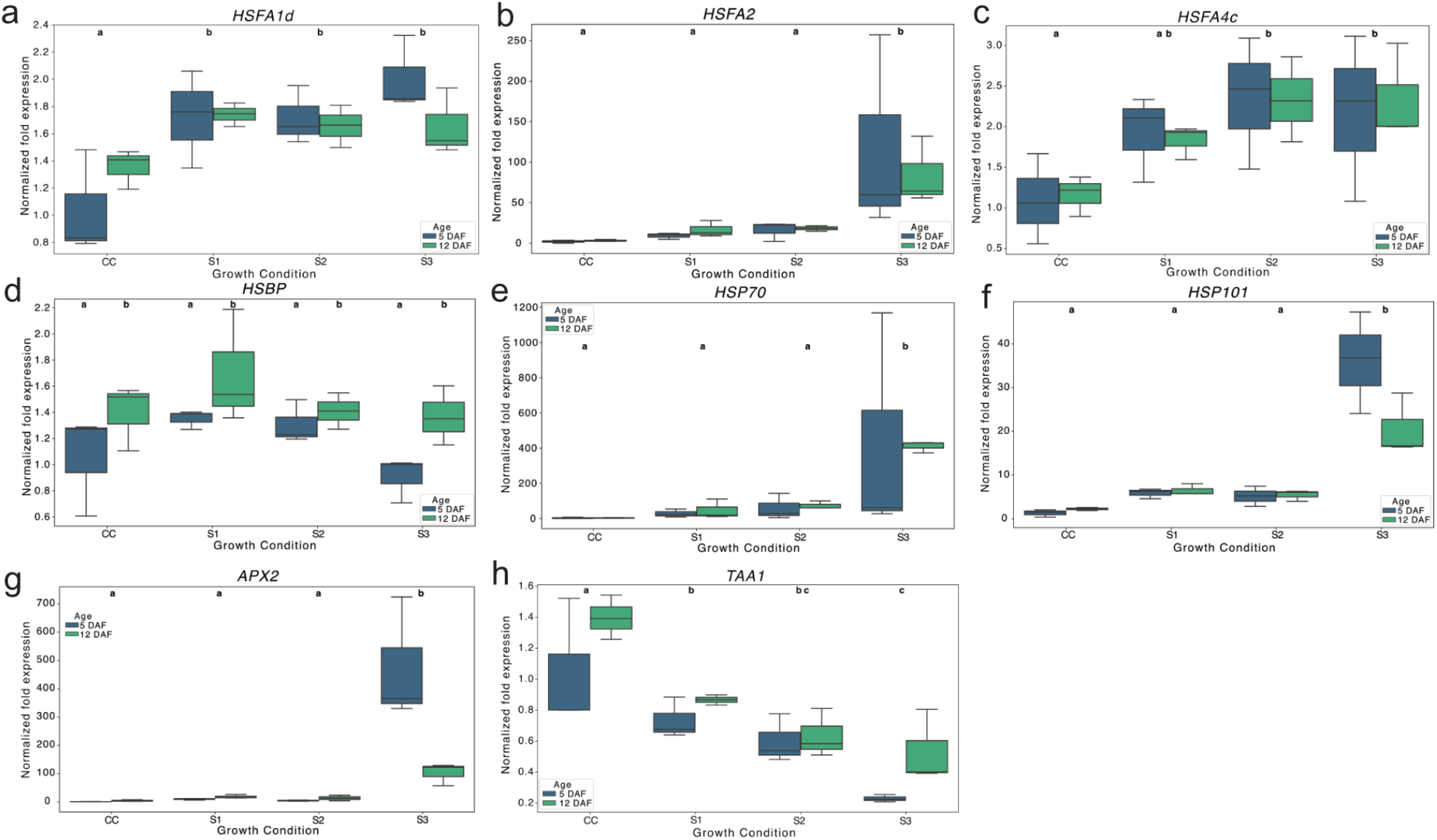
Effect of long-term high temperatures on expression of HSR-related genes in Col-0 pistils. Normalized fold expression assessed by RT-qPCR of *HSFA1d* (**a**), *HSFA2* (b), *HSFA4c* (**c**), *HSBP* (**d**), *HSP70* (**e**), *HSP101* (**f**), *APX2* (**g**), and *TAA1* (**h**) in pistils collected at 5 (blue) and 12 (green) DAF from plant grown under the four growth conditions (CC, S1, S2, S3). Expression levels are normalized to the expression in the 5 DAF pistils at CC. Statistical analysis was performed by two-way ANOVA followed by Tukey HSD tests, showing significant differences between groups with at least *p* < 0.05 to *p* < 0.0001. Groups with the same letter are not significant. CC: Control Condition (21/18 °C); S1: Stress Condition 1 (28/18 °C); S2: Stress Condition 2 (28/24 °C); S3: Stress Condition 3 (34/18 °C).

### The *hot1-3* and *hsbp-2* mutant pollen is defective

The expression analysis indicates that S3 most severely affected the development of reproductive organs. The *hot1-3* and *hspb-2* mutants, which are impaired in HSP101 and HSBP, respectively, are known for their heat-sensitivity (Hong and Vierling 2001; Hsu et al. 2010) and reproductive issues in non-stress conditions (Fu et al. 2002; Qin et al. 2021). We phenotyped the mature pollen of both mutants under CC, mild stress (S1), and severe stress (S3). Pollen development was compromised in both mutant lines. The pollen defects and viability frequencies were similar in the *hsbp-2* mutant and the wild type in CC. Still, S3 caused complete sterility in the *hsbp-2* pollen at 5 DAF (Fig.4c). Partial recovery was observed at 12 DAF with 70 % viable *hsbp-2* pollen (Fig. 4d). On the other hand, 10% of *hot1-3* pollen grains were aborted in CC, reaching 32 % at 5 DAF and 57 % at 12 DAF under S3, indicating an absence of acclimatization (Fig., 4, c and d).

**Figure 4.**
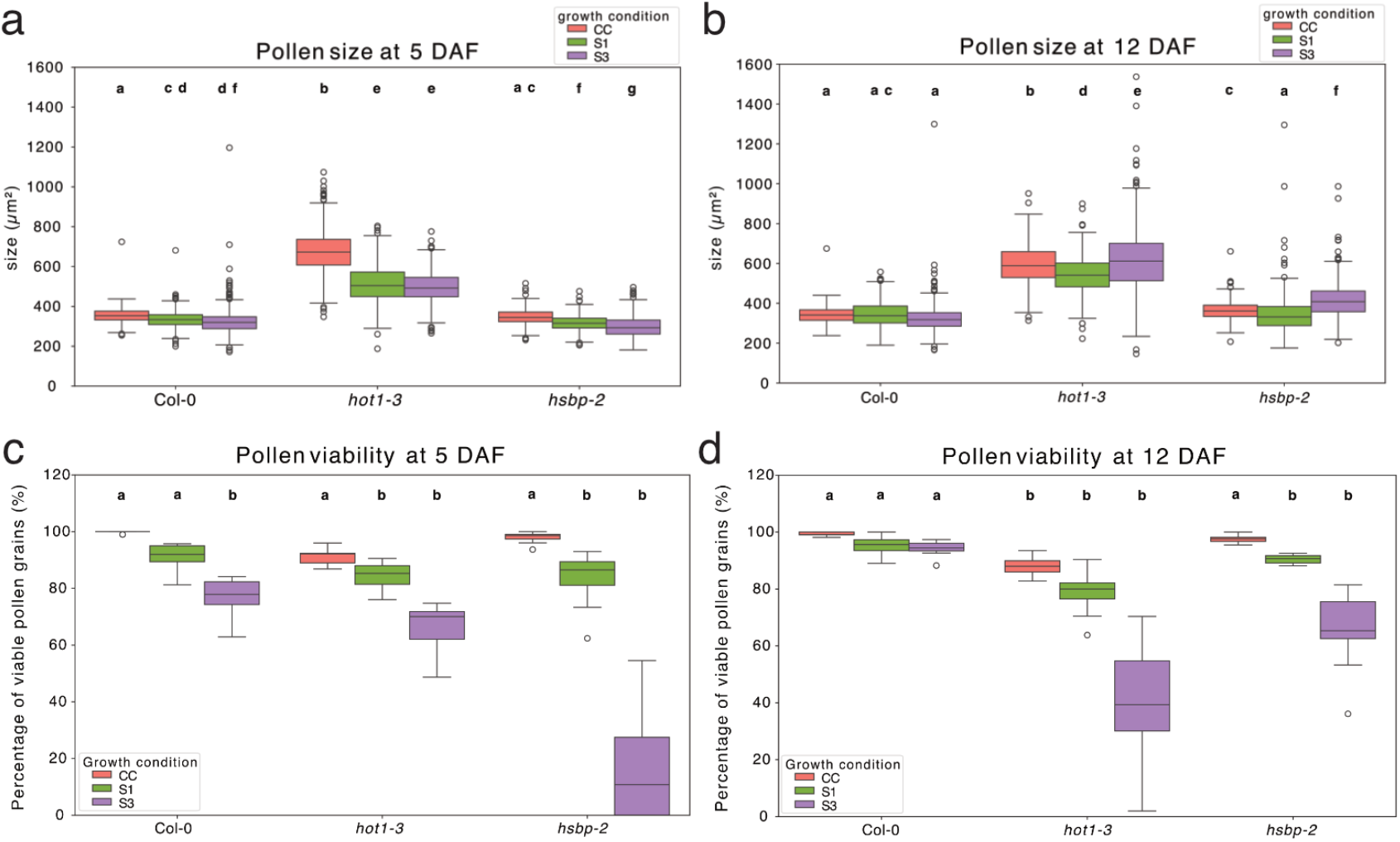
Effect of long-term high temperatures on pollen development in *hot1-3* and *hsbp-2* mutant. (a, b) Pollen size in µm^2^ from 500 grains from 10 plants (Col-0, *hot1-3* and *hsbp-2*) grown under the three growth conditions (CC, control (red); S1, Stress 1 (green); S3, Stress 3 (purple)), assessed at 5 and 12 Days After Flowering (DAF) (n=1874 - 2254). Statistical analysis was performed by two-way ANOVA followed by Tukey HSD tests. (c, d) Pollen viability assessed by counting the number of viable pollen grains from 10 plants grown under the three growth conditions (CC, control (red); S1, Stress 1 (green); S3, Stress 3 (purple)) visualized by Alexander’s stain at 5 and 12 Days After Flowering (DAF). Statistical analysis was performed by Kruskal-Wallis, Mann-Whitney U test and Dunn tests. Data are presented as boxplot with median value, interquartile range as box and minimum and maximum values as whiskers. Groups with the same letter are not significant. CC: Control Condition (21/18 °C); S1: Stress Condition 1 (28/18 °C); S2: Stress Condition 2 (28/24 °C); S3: Stress Condition 3 (34/18 °C).

At 5 DAF, both Col-0 and *hsbp-2* pollen grains reduced their size, with an increased number of outliers for larger and smaller grains in all stress conditions (Fig. 4a). Notably, at 12 DAF, the number of outliers decreased for Col-0 but not in the *hsbp-2* mutant, which produced larger pollen grains at S3 compared to CC (Fig. 4b). The *hot1-3* mutant produced larger pollen grains, twice the size of the Col-0 ones under CC. The overall size of the *hot1-3* pollen grains was reduced at 5 DAF under both stress conditions when compared to CC (Fig. 4a). Under S3 at 12 DAF, while the average size of the *hot1-3* pollen grains was comparable to their size under CC, outliers were twice or three times the standard size (Fig. 4b).

The defects in pollen grains in *hsbp-2* translated into a decreased fertilization rate and seed production (Table 3). In the CC, 39.29 % of ovules were unfertilized (n=2751), and 22.10 % of the developing seeds were aborted in *hsbp-2* mutant, reproducing published data (Hsu et al. 2010). Under stress conditions 1 and 3, 69.62 % and 95.56 % of *hsbp-*2 ovules were unfertilized, producing only 30.38 % and 4.44 % seeds, respectively, among which 20.86 % and 58.54 % aborted. The *hsbp-2* mutants exhibit severe reproductive defects under high-temperature stress. These findings underscore the importance of HSBP and the HSR pathway in maintaining reproductive development under stress conditions.

**Table 3.**
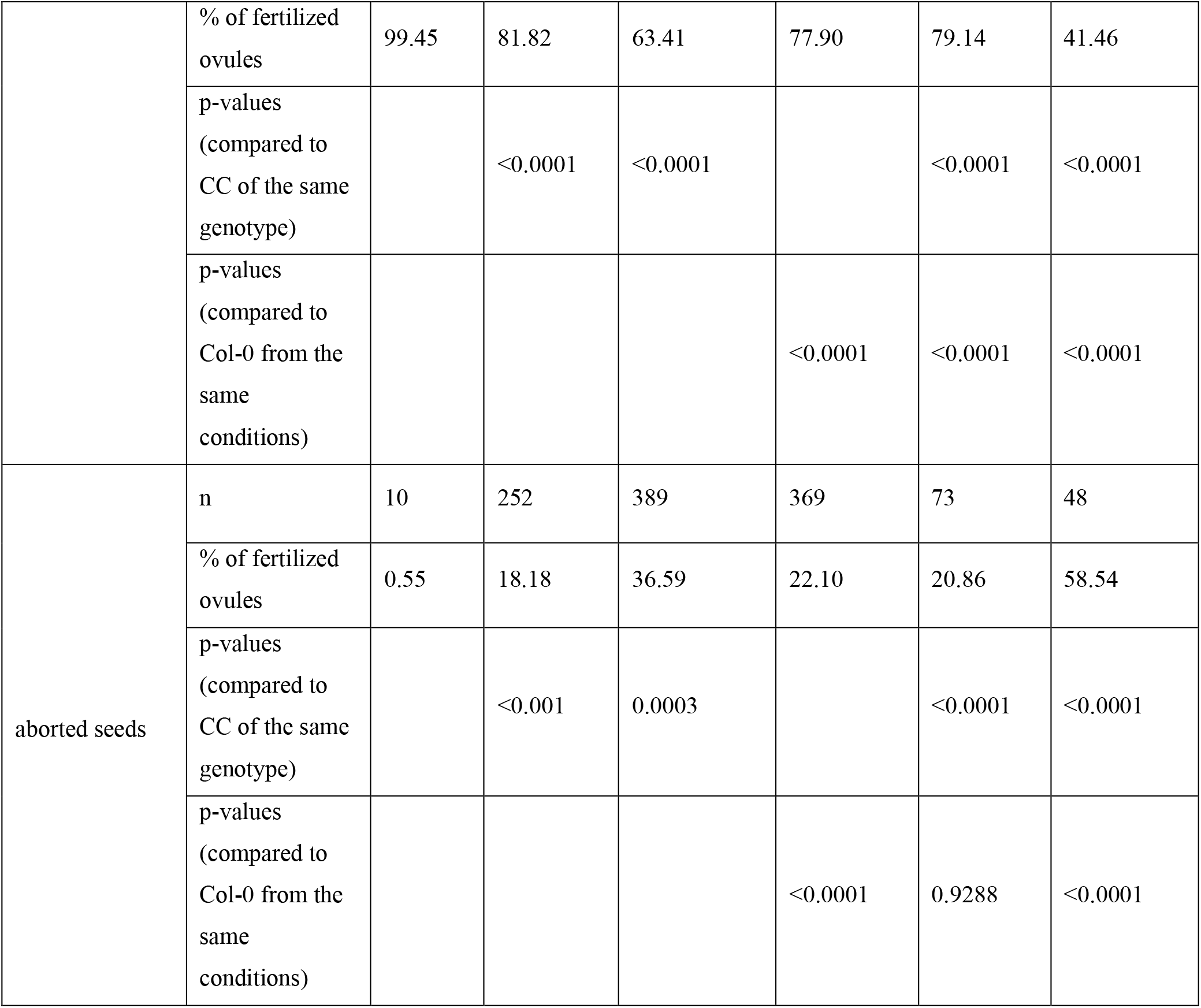
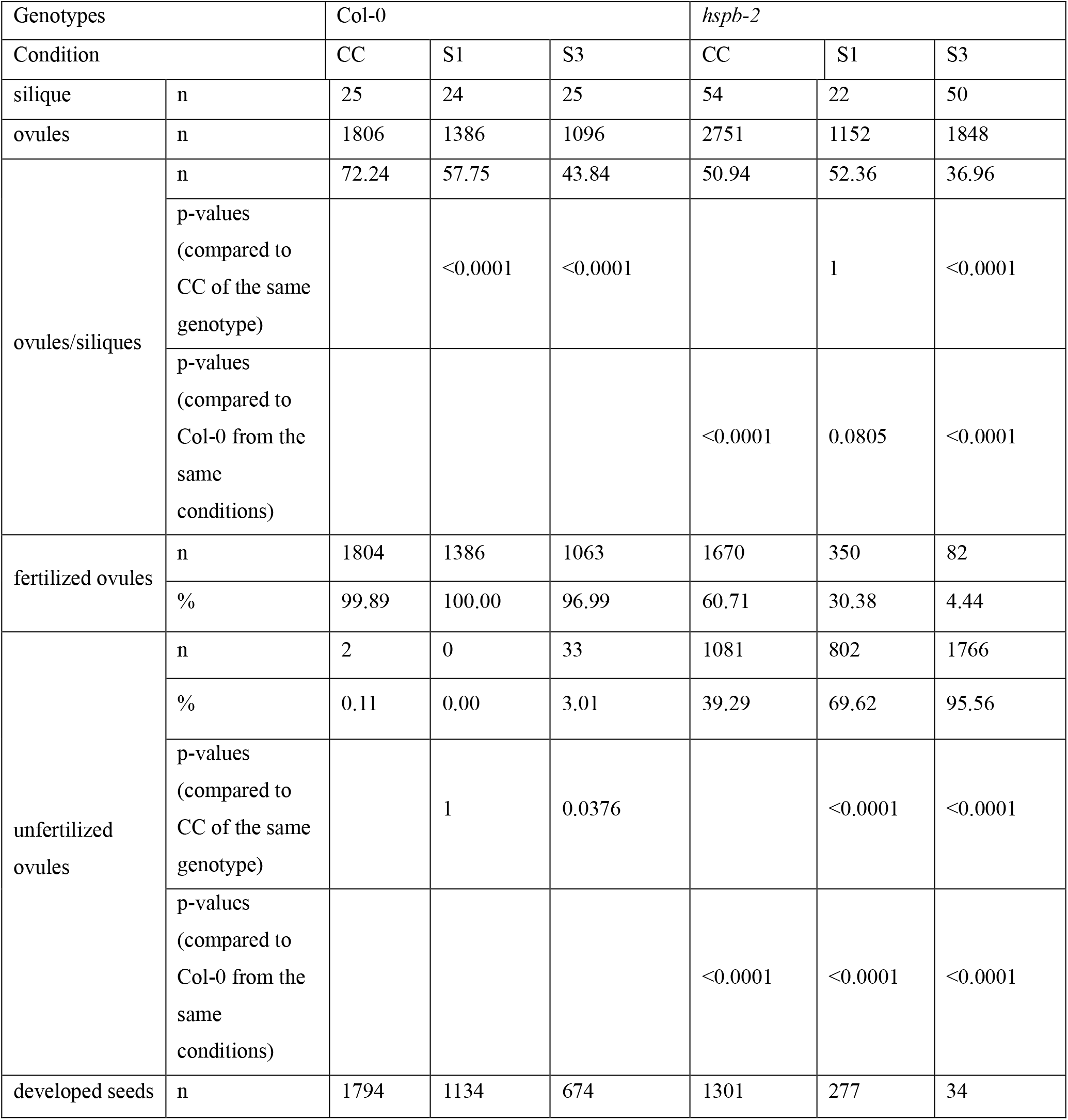
Fertilization rate in Col-0 and *hsbp-2*. Developed seeds, aborted seeds, and unfertilized ovules were counted in n siliques per growth condition. CC, Control Conditions; S1, Stress conditions 1; S3, Stress conditions 3. Statistical analysis was performed Kruskal-Wallis and Dunn multiple comparison tests with *p*-values adjusted by the Holm method. Aborted seeds are also counted as fertilized ovules. Developed seeds are fertilized ovules that form seeds that do not abort.

| Genotypes |  | Col-0 |  |  | *hspb-2* |  |  |
| --- | --- | --- | --- | --- | --- | --- | --- |
| Condition |  | CC | S1 | S3 | CC | S1 | S3 |
| silique | n | 25 | 24 | 25 | 54 | 22 | 50 |
| ovules | n | 1806 | 1386 | 1096 | 2751 | 1152 | 1848 |
| ovules/siliques | n | 72.24 | 57.75 | 43.84 | 50.94 | 52.36 | 36.96 |
|  | p-values<br>(compared to<br>CC of the same<br>genotype) |  | <0.0001 | <0.0001 |  | 1 | <0.0001 |
|  | p-values<br>(compared to<br>Col-0 from the<br>same<br>conditions) |  |  |  | <0.0001 | 0.0805 | <0.0001 |
| fertilized ovules | n | 1804 | 1386 | 1063 | 1670 | 350 | 82 |
|  | % | 99.89 | 100.00 | 96.99 | 60.71 | 30.38 | 4.44 |
| unfertilized<br>ovules | n | 2 | 0 | 33 | 1081 | 802 | 1766 |
|  | % | 0.11 | 0.00 | 3.01 | 39.29 | 69.62 | 95.56 |
|  | p-values<br>(compared to<br>CC of the same<br>genotype) |  | 1 | 0.0376 |  | <0.0001 | <0.0001 |
|  | p-values<br>(compared to<br>Col-0 from the<br>same<br>conditions) |  |  |  | <0.0001 | <0.0001 | <0.0001 |
| developed seeds | n | 1794 | 1134 | 674 | 1301 | 277 | 34 |
|  | % of fertilized ovules | 99.45 | 81.82 | 63.41 | 77.90 | 79.14 | 41.46 |
|  | p-values<br>(compared to CC of the same genotype) |  | <0.0001 | <0.0001 |  | <0.0001 | <0.0001 |
|  | p-values<br>(compared to Col-0 from the same conditions) |  |  |  | <0.0001 | <0.0001 | <0.0001 |
| aborted seeds | n | 10 | 252 | 389 | 369 | 73 | 48 |
|  | % of fertilized ovules | 0.55 | 18.18 | 36.59 | 22.10 | 20.86 | 58.54 |
|  | p-values<br>(compared to CC of the same genotype) |  | <0.001 | 0.0003 |  | <0.0001 | <0.0001 |
|  | p-values<br>(compared to Col-0 from the same conditions) |  |  |  | <0.0001 | 0.9288 | <0.0001 |

### *hot1-3* is tetraploid and can develop aneuploidy

Given the size of *hot1-3* pollen grains, we investigated whether the mutant had issues with mitotic divisions during pollen development. None of the evaluated pollen grains, stained by DAPI, had an abnormal number of nuclei, regardless of the growth condition or size of the pollen grains (Supplemental Videos 1-5). Next, we assessed the ploidy of Col-0 and *hot1-3* sperm nuclei from CC and S3 using flow cytometry. Col-0 consistently showed a haploid DNA cluster at 20 DAPI signal units (Fig. 5a, b), regardless of the temperature regime. However, the *hot1-3* sperm nuclei displayed two clusters corresponding to diploid DNA (50 DAPI signal units) and likely tetraploid DNA (100 DAPI signal units) under S3. This result indicated two unexpected findings: (i) *hot1-3* plants were tetraploid and (ii) produced unreduced microspores (Fig. 5c, d). We confirmed the tetraploidy of *hot1-3* plants in leaf samples (Fig. 5e-h, Supplementary Fig. S3). Furthermore, staining anaphase chromosomes with DAPI identified ten segregating chromosomes in Col-0 sepals, but twenty chromosomes in *hot1-3* sepals (Supplemental videos 6 and 7). Overlaying the chromatographs from leaves revealed a double peak at 200 DAPI signal units and two peaks at 350 and 400 DAPI signal units, identifying that some *hot1-3* plants are also aneuploid (Fig. 5h).

**Figure 5.**
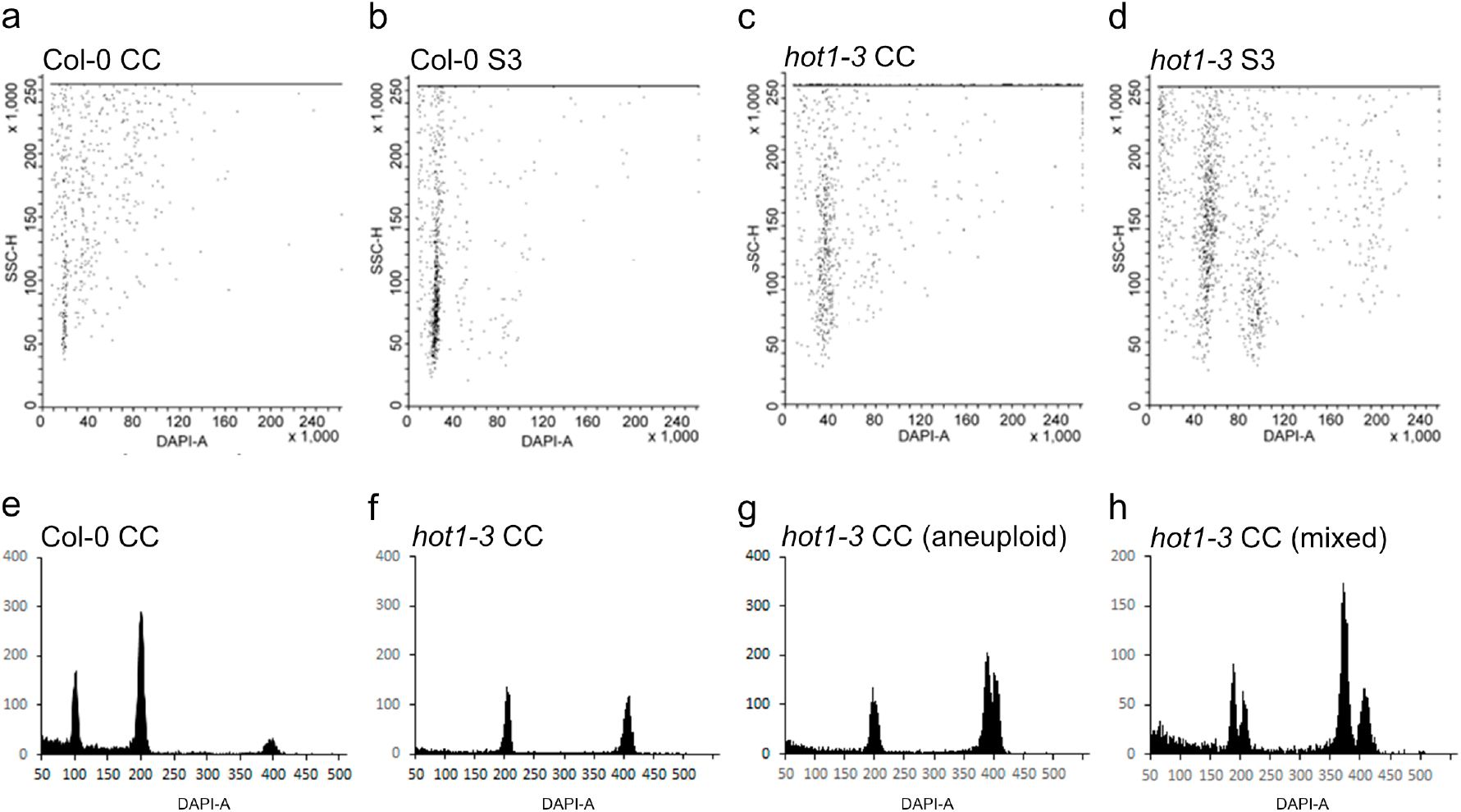
*hot1-3* is tetraploid and aneuploid. (a-d) Bivariate scatter plot of sperm nuclei from Col-0 (a, b) and *hot1-3* mutant (c, d) plants grown under the CC (a, c) and S3 (b, d) growth conditions. The x-axis shows relative DAPI intensity (DNA content) and the y-axis side scatter pulse height (SSC-h), indicating the running time of the nuclei through the sensor. (e-h) Flow cytometry histograms showing representative profiles of peaks from diploid Col-0 (e) and *hot1-3* mutant (f-h) plants grown under the CC growth condition. The first peak corresponds to 2C, the second to 4C and the third to 8C nuclei (8C peak not shown in *hot1-*3). The histograms show two types of ploidy in *hot1-3* mutant: tetraploid (f) and aneuploid (g). The histogram in (h) displays the overlap when the material of the two plants is mixed prior analysis.

To confirm whether the ploidy change was due to the inactivation of HSP101, we assessed the ploidy from the original *hot1-3* seeds obtained from NASC (N16284) and two other *hsp101* mutant alleles (SALK_066374C and SALK_099583C). The original *hot1-3* mutant plants were tetraploid, but the two other *hsp101* mutant lines were diploid (Supplementary Fig. S3), suggesting that the *hot1-3* became tetraploid at some point during the selection and characterisation of the mutant allele (Hong and Vierling 2001).

## Discussion

In recent years, the global climate crisis has significantly reduced crop production (Sidhu, 2023), highlighting the need to understand plant adaptive strategies in response to high temperatures. To address this, we studied different high-temperature scenarios using *Arabidopsis thaliana* (this work) and *Brassica napus* (Mácová et al. 2022) as model plants. Previous research primarily focused on plant reactions to short heat stress, which is less representative of natural conditions. In our study, we designed different nature-like scenarios that allow plants to grow and adapt to different heat stress conditions, providing comprehensive insights into plant thermo-adaptation to high-temperature environments.

Our study found that flowering in Arabidopsis at 34 °C results in a reduced number of fertile flowers and decreased fertility. Previous studies have linked this reduction in crop production to heat-induced pollen sterility (Endo et al. 2009; Begcy et al. 2019). While high temperatures did affect pollen development in our growth setup, it adapted to long-term heat stress, likely due to cooler night temperatures. Unlike our approach, most Arabidopsis studies use intense heat shocks (Lei et al. 2020; Rutley et al. 2021) or constant high temperatures over long periods (Yoshida et al. 2008; Hedhly et al. 2020), conditions that are uncommon in nature and cause severe plant damages. High temperatures also affect the female gametophyte in other plant species (Wang et al. 2021; Shi et al. 2022; Jedličková et al. 2023), but this process is not understood in *Arabidopsis thaliana*. Consequently, damage to both gametophytes may result in reduced pollination and fertilization rates, significantly decreasing seed production (Jiang et al., 2018).

Expression profiling showed that 34 °C severely affects plant metabolism. Stress-related genes, such as *HSP101, HSP70*, and *APX2*, were extremely upregulated in pistils of plants grown under stress conditions, while transcription factors (*HSFA2, HSFA1d*, or *HSFA4c*) regulating HSR were upregulated to a lesser extent. Chaperones like HSP101 are key to thermo-adaptation (Gurley 2000; Queitsch et al. 2000; Hong and Vierling 2001; Nieto-Sotelo et al. 2002; Wu et al. 2013). Adverse environments induce Reactive Oxygen Species (ROS) production, such as H_2_O_2_, which must be detoxified, notably by peroxidases, for example during abiotic stresses (Chang et al. 2009), as well as during developmental processes (Ye et al. 2000; Jacobowitz et al. 2019).

The Arabidopsis *hot1-3* mutant, characterized by a T-DNA insertion in the second exon of the *HSP101* gene (Hong and Vierling 2001), displays severe defects during male gametophyte development when grown under high temperature conditions. This underscores the critical role of HSP101 in protecting gametes during thermal stress. Optimal temperatures are essential for meiotic crossovers (Lloyd et al. 2018) and chromosome segregation (Lei et al. 2020; Khaitova et al. 2023). Our study revealed that the *hot1-3* mutant is tetraploid. This line has a complex origin and, therefore, it is unclear when the polyploidization occurred. However, it was likely during early generation as it is already present in the material donated to the stock centre by Suk-Whan Hong (Hong and Vierling 2001). Interestingly, the *hot1-3* can also develop aneuploidy, promoted by high temperatures. Heat stress is known to induce unreduced microspores (Storme and Geelen 2020), as observed in Arabidopsis Col-0 plants grown at 35/22 °C (Nguyen et al. 2019). However, this effect was not seen under our Stress Condition 3, which could be attributed to the different temperature fluctuations between day and night (13 °C in Nguyen et al. (2019) versus 18 °C in our study). HSFs coordinate both the HSR and plant development. HSBP, a negative regulator of HSF transcriptional activity, attenuates the HSR when temperatures decrease (Hsu et al. 2010). Using the *hsbp-2* mutant, which cannot efficiently attenuate the HSR, we examined how Arabidopsis seeds develop under our stress conditions. HSBP regulates seed development (Hsu et al. 2010) and meiotic crossover during ovule development (Kim et al. 2022). Our findings show that *hsbp-2* pollen development is susceptible to high temperatures. The analysis of a hypersensitive mutant to temperature stress (*hot1-3*) and a hyperreactive mutant (*hsbp-2*) provided insights into the reproductive phase under temperature stress.

## Data availability

Source data are provided in the Zenodo repository (https://doi.org/10.5281/zenodo.19607439).

## Author Contributions

J.F.S.L., A.P., and H.S.R. designed the research; J.F.S.L., M.Š, and F.Y. performed the analysis and analyzed the data; J.F.S.L. and H.S.R. wrote the article. All authors read and approved the manuscript.

## Acknowledgments

We would like to thank E. Vierling (University of Massachusetts Amherst, USA) for the *hot1-3* seeds, and NASC for providing the remaining lines. We acknowledge the Core Facility CELLIM supported by the Czech-BioImaging large RI project (LM2023050 funded by MEYS CR) and the Core Facility Plant Sciences of CEITEC Masaryk University for their support in obtaining scientific data presented in this paper.

## References

Agho C, Avni A, Bacu A, et al (2024) Integrative approaches to enhance reproductive resilience of crops for climate-proof agriculture. Plant Stress 100704. 10.1016/j.stress.2024.100704

Ai H, Bellstaedt J, Bartusch KS, et al (2023) Auxin-dependent regulation of cell division rates governs root thermomorphogenesis. EMBO J 42:e111926. 10.15252/embj.2022111926

Alexander MP (1969) Differential staining of aborted and nonaborted pollen. Stain technology 44:117– 122. 10.3109/10520296909063335

Andersen CL, Jensen JL, Ørntoft TF (2004) Normalization of real-time quantitative reverse transcription-PCR data: a model-based variance estimation approach to identify genes suited for normalization, applied to bladder and colon cancer data sets. Cancer research 64:5245–5250. 10.1158/0008-5472.can-04-0496

Andrási N, Rigó G, Zsigmond L, et al (2019) The mitogen-activated protein kinase 4-phosphorylated heat shock factor A4A regulates responses to combined salt and heat stresses. J Exp Bot 70:4903–4918. 10.1093/jxb/erz217

Avin-Wittenberg T (2018) Autophagy and its role in plant abiotic stress management. Plant, Cell Environ 42:1045–1053. 10.1111/pce.13404

Barnabás B, Jäger K, Fehér A (2008) The effect of drought and heat stress on reproductive processes in cereals. Plant, Cell & Environment 31:11–38. 10.1111/j.1365-3040.2007.01727.x

Begcy K, Nosenko T, Zhou L-Z, et al (2019) Male Sterility in Maize after Transient Heat Stress during the Tetrad Stage of Pollen Development. Plant Physiol 181:683–700. 10.1104/pp.19.00707

Borges F, Gardner R, Lopes T, et al (2012) FACS-based purification of Arabidopsis microspores, sperm cells and vegetative nuclei. Plant Methods 8:44. 10.1186/1746-4811-8-44

Boyes DC, Zayed AM, Ascenzi R, et al (2001) Growth Stage–Based Phenotypic Analysis of Arabidopsis. Plant Cell 13:1499–1510. 10.1105/tpc.010011

Carranco R, Prieto-Dapena P, Almoguera C, Jordano J (2017) SUMO-Dependent Synergism Involving Heat Shock Transcription Factors with Functions Linked to Seed Longevity and Desiccation Tolerance. Front Plant Sci 8:974. 10.3389/fpls.2017.00974

Chang CCC, Ślesak I, Jordá L, et al (2009) Arabidopsis Chloroplastic Glutathione Peroxidases Play a Role in Cross Talk between Photooxidative Stress and Immune Responses. Plant Physiol 150:670– 683. 10.1104/pp.109.135566

Chan-Schaminet KY, Baniwal SK, Bublak D, et al (2009) Specific Interaction between Tomato HsfA1 and HsfA2 Creates Hetero-oligomeric Superactivator Complexes for Synergistic Activation of Heat Stress Gene Expression. Journal of Biological Chemistry 284:20848–20857. 10.1074/jbc.m109.007336

Charng Y, Liu H, Liu N, et al (2007) A Heat-Inducible Transcription Factor, HsfA2, Is Required for Extension of Acquired Thermotolerance in Arabidopsis. Plant Physiol 143:251–262. 10.1104/pp.106.091322

Cohen-Peer R, Schuster S, Meiri D, et al (2010) Sumoylation of Arabidopsis heat shock factor A2 (HsfA2) modifies its activity during acquired thermotholerance. Plant Mol Biol 74:33–45. 10.1007/s11103-010-9652-1

Deng Y, Srivastava R, Quilichini TD, et al (2016) IRE1, a component of the unfolded protein response signaling pathway, protects pollen development in Arabidopsis from heat stress. Plant J 88:193–204. 10.1111/tpj.13239

Endo M, Tsuchiya T, Hamada K, et al (2009) High Temperatures Cause Male Sterility in Rice Plants with Transcriptional Alterations During Pollen Development. Plant Cell Physiol 50:1911–1922. 10.1093/pcp/pcp135

Evrard A, Kumar M, Lecourieux D, et al (2013) Regulation of the heat stress response in Arabidopsis by MPK6-targeted phosphorylation of the heat stress factor HsfA2. PeerJ 1:e59. 10.7717/peerj.59

Friedrich T, Oberkofler V, Trindade I, et al (2021) Heteromeric HSFA2/HSFA3 complexes drive transcriptional memory after heat stress in Arabidopsis. Nat Commun 12:3426. 10.1038/s41467-021-23786-6

Fu S, Meeley R, Scanlon MJ (2002) Empty pericarp2 encodes a negative regulator of the heat shock response and is required for maize embryogenesis. Plant cell 14:3119–32. 10.1105/tpc.006726

Fu S, Rogowsky P, Nover L, Scanlon MJ (2005) The maize heat shock factor-binding protein paralogs EMP2 and HSBP2 interact non-redundantly with specific heat shock factors. Planta 224:42–52. 10.1007/s00425-005-0191-y

Gurley WB (2000) HSP101: A Key Component for the Acquisition of Thermotolerance in Plants. Plant Cell 12:457. 10.2307/3871060

Hahn A, Bublak D, Schleiff E, Scharf K-D (2011) Crosstalk between Hsp90 and Hsp70 Chaperones and Heat Stress Transcription Factors in Tomato. Plant Cell 23:741–755. 10.1105/tpc.110.076018

Hedhly A, Nestorova A, Herrmann A, Grossniklaus U (2020) Acute heat stress during stamen development affects both the germline and sporophytic lineages in Arabidopsis thaliana (L.) Heynh. Environmental and Experimental Botany 173:103992. 10.1016/j.envexpbot.2020.103992

Hong SW, Vierling E (2001) Hsp101 is necessary for heat tolerance but dispensable for development and germination in the absence of stress. Plant J Cell Mol Biology 27:25–35

Hsu S-F, Lai H-C, Jinn T-L (2010) Cytosol-Localized Heat Shock Factor-Binding Protein, AtHSBP, Functions as a Negative Regulator of Heat Shock Response by Translocation to the Nucleus and Is Required for Seed Development in Arabidopsis. Plant Physiol 153:773–784. 10.1104/pp.109.151225

Huang Y-C, Liu C-C, Li Y-J, et al (2024) Multifaceted roles of Arabidopsis heat shock factor binding protein in plant growth, development, and heat shock response. Environ Exp Bot 226:105878. 10.1016/j.envexpbot.2024.105878

Jacob P, Hirt H, Bendahmane A (2017) The heat-shock protein/chaperone network and multiple stress resistance. Plant Biotechnology Journal 15:405–414. 10.1111/pbi.12659

Jacobowitz JR, Doyle WC, Weng J-K (2019) PRX9 and PRX40 Are Extensin Peroxidases Essential for Maintaining Tapetum and Microspore Cell Wall Integrity during Arabidopsis Anther Development. Plant Cell 31:848–861. 10.1105/tpc.18.00907

Jedličková V, Hejret V, Demko M, et al (2023) Transcriptome analysis of thermomorphogenesis in ovules and during early seed development in Brassica napus. BMC Genomics 24:236. 10.1186/s12864-023-09316-2

Jegadeesan S, Beery A, Altahan L, et al (2018) Ethylene production and signaling in tomato (Solanum lycopersicum) pollen grains is responsive to heat stress conditions. Plant Reprod 31:367–383. 10.1007/s00497-018-0339-0

Khaitova LC, Mikulkova P, Pecinkova J, et al (2024) Heat stress impairs centromere structure and segregation of meiotic chromosomes in Arabidopsis. eLife 12:RP90253. 10.7554/elife.90253

Khaitova LC, Mikulkova P, Pecinkova J, et al (2023) Heat stress impairs centromere structure and segregation of meiotic chromosomes in Arabidopsis. 10.7554/elife.90253.1

Khan AH, Min L, Ma Y, et al (2023) High-temperature stress in crops: male sterility, yield loss and potential remedy approaches. Plant Biotechnol J 21:680–697. 10.1111/pbi.13946

Kim J, Park J, Kim H, et al (2022) Arabidopsis HEAT SHOCK FACTOR BINDING PROTEIN is required to limit meiotic crossovers and HEI10 transcription. EMBO J 41:e109958. 10.15252/embj.2021109958

Kim M, Lee U, Small I, et al (2012) Mutations in an Arabidopsis Mitochondrial Transcription Termination Factor–Related Protein Enhance Thermotolerance in the Absence of the Major Molecular Chaperone HSP101. Plant Cell 24:3349–3365. 10.1105/tpc.112.101006

Koini MA, Alvey L, Allen T, et al (2009) High Temperature-Mediated Adaptations in Plant Architecture Require the bHLH Transcription Factor PIF4. Current biology 19:408–413. 10.1016/j.cub.2009.01.046

Lei X, Ning Y, Elesawi IE, et al (2020) Heat stress interferes with chromosome segregation and cytokinesis during male meiosis in Arabidopsis thaliana. Plant Signal Behav 15:1746985. 10.1080/15592324.2020.1746985

Lin J-S, Kuo C-C, Yang I-C, et al (2018a) MicroRNA160 Modulates Plant Development and Heat Shock Protein Gene Expression to Mediate Heat Tolerance in Arabidopsis. Front Plant Sci 9:68. 10.3389/fpls.2018.00068

Lin K-F, Tsai M-Y, Lu C-A, et al (2018b) The roles of Arabidopsis HSFA2, HSFA4a, and HSFA7a in the heat shock response and cytosolic protein response. Bot Stud 59:15. 10.1186/s40529-018-0231-0

Lloyd A, Morgan C, Franklin FCH, Bomblies K (2018) Plasticity of Meiotic Recombination Rates in Response to Temperature in Arabidopsis. Genetics 208:1409–1420. 10.1534/genetics.117.300588

Luo J, Jiang J, Sun S, Wang X (2022) Brassinosteroids promote thermotolerance through releasing BIN2-mediated phosphorylation and suppression of HsfA1 transcription factors in Arabidopsis. Plant Commun 3:100419. 10.1016/j.xplc.2022.100419

Mácová K, Prabhullachandran U, Štefková M, et al (2022) Long-Term High-Temperature Stress Impacts on Embryo and Seed Development in Brassica napus. Front Plant Sci 13:844292. 10.3389/fpls.2022.844292

McLoughlin F, Kim M, Marshall RS, et al (2019) HSP101 Interacts with the Proteasome and Promotes the Clearance of Ubiquitylated Protein Aggregates. PLANT PHYSIOLOGY pp.00263.2019-52. 10.1104/pp.19.00263

Merret R, Carpentier M-C, Favory J-J, et al (2017) Heat Shock Protein HSP101 Affects the Release of Ribosomal Protein mRNAs for Recovery after Heat Shock. Plant Physiol 174:1216–1225. 10.1104/pp.17.00269

Mizoi J, Kanazawa N, Kidokoro S, et al (2019) Heat-induced inhibition of phosphorylation of the stress-protective transcription factor DREB2A promotes thermotolerance of Arabidopsis thaliana. J Biol Chem 294:902–917. 10.1074/jbc.ra118.002662

Morimoto RI (1998) Regulation of the heat shock transcriptional response: cross talk between a family of heat shock factors, molecular chaperones, and negative regulators. Genes Dev 12:3788–3796. 10.1101/gad.12.24.3788

Muthusamy M, Son S, Park SR, Lee SI (2023) Heat shock factor binding protein BrHSBP1 regulates seed and pod development in Brassica rapa. Front Plant Sci 14:1232736. 10.3389/fpls.2023.1232736

Nguyen TD, Jang S, Soh M-S, et al (2019) High daytime temperature induces male sterility with developmental defects in male reproductive organs of Arabidopsis. Plant Biotechnol Rep 13:635–643. 10.1007/s11816-019-00559-8

Nieto-Sotelo J, Martínez LM, Ponce G, et al (2002) Maize HSP101 Plays Important Roles in Both Induced and Basal Thermotolerance and Primary Root Growth. Plant Cell 14:1621–1633. 10.1105/tpc.010487

Nover L, Bharti K, Dring P, et al (2001) Arabidopsis and the heat stress transcription factor world: how many heat stress transcription factors do we need? Cell Stress Chaperones 6:177–189. 10.1379/1466-1268(2001)006<0177:aathst>2.0.co;2

Ogawa D, Yamaguchi K, Nishiuchi T (2007) High-level overexpression of the Arabidopsis HsfA2 gene confers not only increased themotolerance but also salt/osmotic stress tolerance and enhanced callus growth. J Exp Bot 58:3373–3383. 10.1093/jxb/erm184

Prerostova S, Dobrev PI, Kramna B, et al (2020) Heat Acclimation and Inhibition of Cytokinin Degradation Positively Affect Heat Stress Tolerance of Arabidopsis. Front Plant Sci 11:87. 10.3389/fpls.2020.00087

Qin F, Yu B, Li W (2021) Heat shock protein 101 (HSP101) promotes flowering under nonstress conditions. Plant Physiol 186:407–419. 10.1093/plphys/kiab052

Queitsch C, Hong S-W, Vierling E, Lindquist S (2000) Heat Shock Protein 101 Plays a Crucial Role in Thermotolerance in Arabidopsis. Plant Cell 12:479. 10.2307/3871063

Raja MM, Vijayalakshmi G, Naik ML, et al (2019) Pollen development and function under heat stress: from effects to responses. Acta Physiol Plant 41:47. 10.1007/s11738-019-2835-8

Rana RM, Dong S, Tang H, et al (2012) Functional analysis of OsHSBP1 and OsHSBP2 revealed their involvement in the heat shock response in rice (Oryza sativa L.). J Exp Bot 63:6003–6016. 10.1093/jxb/ers245

Rieu I, Twell D, Firon N (2017) Pollen Development at High Temperature: from Acclimation to Collapse. PLANT PHYSIOLOGY. 10.1104/pp.16.01644

Rutley N, Poidevin L, Doniger T, et al (2021) Characterization of novel pollen-expressed transcripts reveals their potential roles in pollen heat stress response in Arabidopsis thaliana. Plant Reprod 34:61– 78. 10.1007/s00497-020-00400-1

Sage TL, Bagha S, Lundsgaard-Nielsen V, et al (2015) The effect of high temperature stress on male and female reproduction in plants. Field Crops Research 182:30–42. 10.1016/j.fcr.2015.06.011

Satyal SH, Chen D, Fox SG, et al (1998) Negative regulation of the heat shock transcriptional response by HSBP1. Genes Dev 12:1962–1974. 10.1101/gad.12.13.1962

Scharf K-D, Berberich T, Ebersberger I, Nover L (2012) The plant heat stress transcription factor (Hsf) family: structure, function and evolution. Biochimica et biophysica acta 1819:104–119. 10.1016/j.bbagrm.2011.10.002

Shi W, Yang J, Kumar R, et al (2022) Heat Stress During Gametogenesis Irreversibly Damages Female Reproductive Organ in Rice. Rice 15:32. 10.1186/s12284-022-00578-0

Shi Y, Mosser DD, Morimoto RI (1998) Molecularchaperones as HSF1-specific transcriptional repressors. Gene Dev 12:654–666. 10.1101/gad.12.5.654

Snider JL, Oosterhuis DM, Loka DA, Kawakami EM (2011) High temperature limits in vivo pollen tube growth rates by altering diurnal carbohydrate balance in field-grown Gossypium hirsutum pistils. Journal of Plant Physiology 168:1168–1175. 10.1016/j.jplph.2010.12.011

Song C, Lee J, Kim T, et al (2018) VOZ1, a transcriptional repressor of DREB2C, mediates heat stress responses in Arabidopsis. Planta 247:1439–1448. 10.1007/s00425-018-2879-9

Storme ND, Geelen D (2020) High temperatures alter cross-over distribution and induce male meiotic restitution in Arabidopsis thaliana. Commun Biol 3:187. 10.1038/s42003-020-0897-1

Suzuki N, Bassil E, Hamilton JS, et al (2016) ABA Is Required for Plant Acclimation to a Combination of Salt and Heat Stress. PLoS ONE 11:e0147625. 10.1371/journal.pone.0147625

Tak Y, Lal SS, Gopan S, et al (2022) Identification of subfunctionalized aggregate-remodeling J-domain proteins in Arabidopsis thaliana. J Exp Bot 74:1705–1722. 10.1093/jxb/erac514

Taylor SC, Nadeau K, Abbasi M, et al (2019) The Ultimate qPCR Experiment: Producing Publication Quality, Reproducible Data the First Time. Trends Biotechnol 37:761–774. 10.1016/j.tibtech.2018.12.002

Teng C, Dong H, Shi L, et al (2008) Serine palmitoyltransferase, a key enzyme for de novo synthesis of sphingolipids, is essential for male gametophyte development in Arabidopsis. Plant Physiol 146:1322– 32. 10.1104/pp.107.113506

Vacca RA, Pinto MC de, Valenti D, et al (2004) Production of Reactive Oxygen Species, Alteration of Cytosolic Ascorbate Peroxidase, and Impairment of Mitochondrial Metabolism Are Early Events in Heat Shock-Induced Programmed Cell Death in Tobacco Bright-Yellow 2 Cells. Plant Physiol 134:1100–1112. 10.1104/pp.103.035956

Vandesompele J, Preter KD, Pattyn F, et al (2002) Accurate normalization of real-time quantitative RT-PCR data by geometric averaging of multiple internal control genes. Genome Biol 3:research0034.1. 10.1186/gb-2002-3-7-research0034

Wahid A, Gelani S, Ashraf M, Foolad MR (2007) Heat tolerance in plants: An overview. Environ Exp Bot 61:199–223. 10.1016/j.envexpbot.2007.05.011

Wang T-Y, Wu J-R, Duong NKT, et al (2021) HSP70-4 and farnesylated AtJ3 constitute a specific HSP70/HSP40-based chaperone machinery essential for prolonged heat stress tolerance in Arabidopsis. J Plant Physiol 261:153430. 10.1016/j.jplph.2021.153430

Wu T, Juan Y, Hsu Y, et al (2013) Interplay between Heat Shock Proteins HSP101 and HSA32 Prolongs Heat Acclimation Memory Posttranscriptionally in Arabidopsis. Plant Physiol 161:2075–2084. 10.1104/pp.112.212589

Yang D, Li Y, Shi Y, et al (2016) Exogenous Cytokinins Increase Grain Yield of Winter Wheat Cultivars by Improving Stay-Green Characteristics under Heat Stress. PLoS ONE 11:e0155437. 10.1371/journal.pone.0155437

Yang F, Fernández-Jiménez N, Tučková M, et al (2021) Defects in meiotic chromosome segregation lead to unreduced male gametes in Arabidopsis SMC5/6 complex mutants. Plant cell 33:3104–3119. 10.1093/plcell/koab178

Ye, Rodriguez, Tran, et al (2000) The developmental transition to flowering represses ascorbate peroxidase activity and induces enzymatic lipid peroxidation in leaf tissue in Arabidopsis thaliana. Plant Sci : Int J Exp plant Biol 158:115–127. 10.1016/s0168-9452(00)00316-2

Yoshida T, Sakuma Y, Todaka D, et al (2008) Functional analysis of an Arabidopsis heat-shock transcription factor HsfA3 in the transcriptional cascade downstream of the DREB2A stress-regulatory system. Biochem Biophys Res Commun 368:515–521. 10.1016/j.bbrc.2008.01.134

Young LW, Wilen RW, Bonham-Smith PC (2004) High temperature stress of Brassica napus during flowering reduces micro- and megagametophyte fertility, induces fruit abortion, and disrupts seed production. Journal of Experimental Botany 55:485–495. 10.1093/jxb/erh038

Zhang C, Li G, Chen T, et al (2018) Heat stress induces spikelet sterility in rice at anthesis through inhibition of pollen tube elongation interfering with auxin homeostasis in pollinated pistils. Rice 11:14. 10.1186/s12284-018-0206-5

Zinn KE, Tunc-Ozdemir M, Harper JF (2010) Temperature stress and plant sexual reproduction: uncovering the weakest links. Journal of Experimental Botany 61:1959–1968. 10.1093/jxb/erq053

Zupanska AK, LeFrois C, Ferl RJ, Paul A-L (2019) HSFA2 Functions in the Physiological Adaptation of Undifferentiated Plant Cells to Spaceflight. Int J Mol Sci 20:390. 10.3390/ijms20020390

