## Supplementary Figures for "Adaptive response to long-term high temperatures during the reproductive development in *Arabidopsis thaliana*"

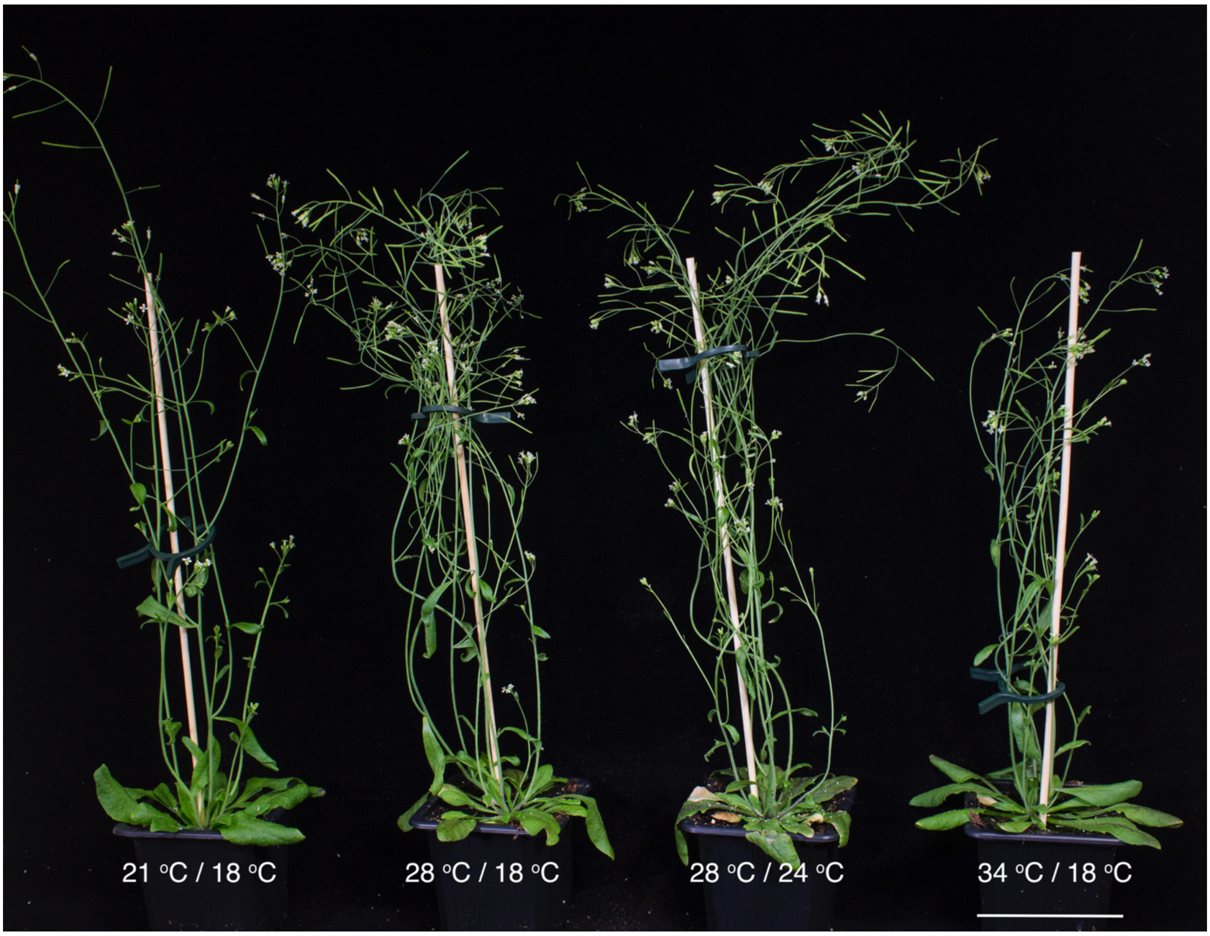


**Supplementary Figure S1.** Phenotypes of flowering plants, two weeks after transfer to the indicated growth temperatures.

Columbia plants are grown under Control Condition (21 °C / 18 °C) until the initiation of flowering. Plants are kept in the same condition or transferred to Stress Condition S1 (28 °C / 18 °C), Stress Condition S2 (28 °C / 24 °C), or Stress Condition S3 (34 °C / 18 °C). The picture was taken 14 days after the transfer. Scale bar represents 6 cm.

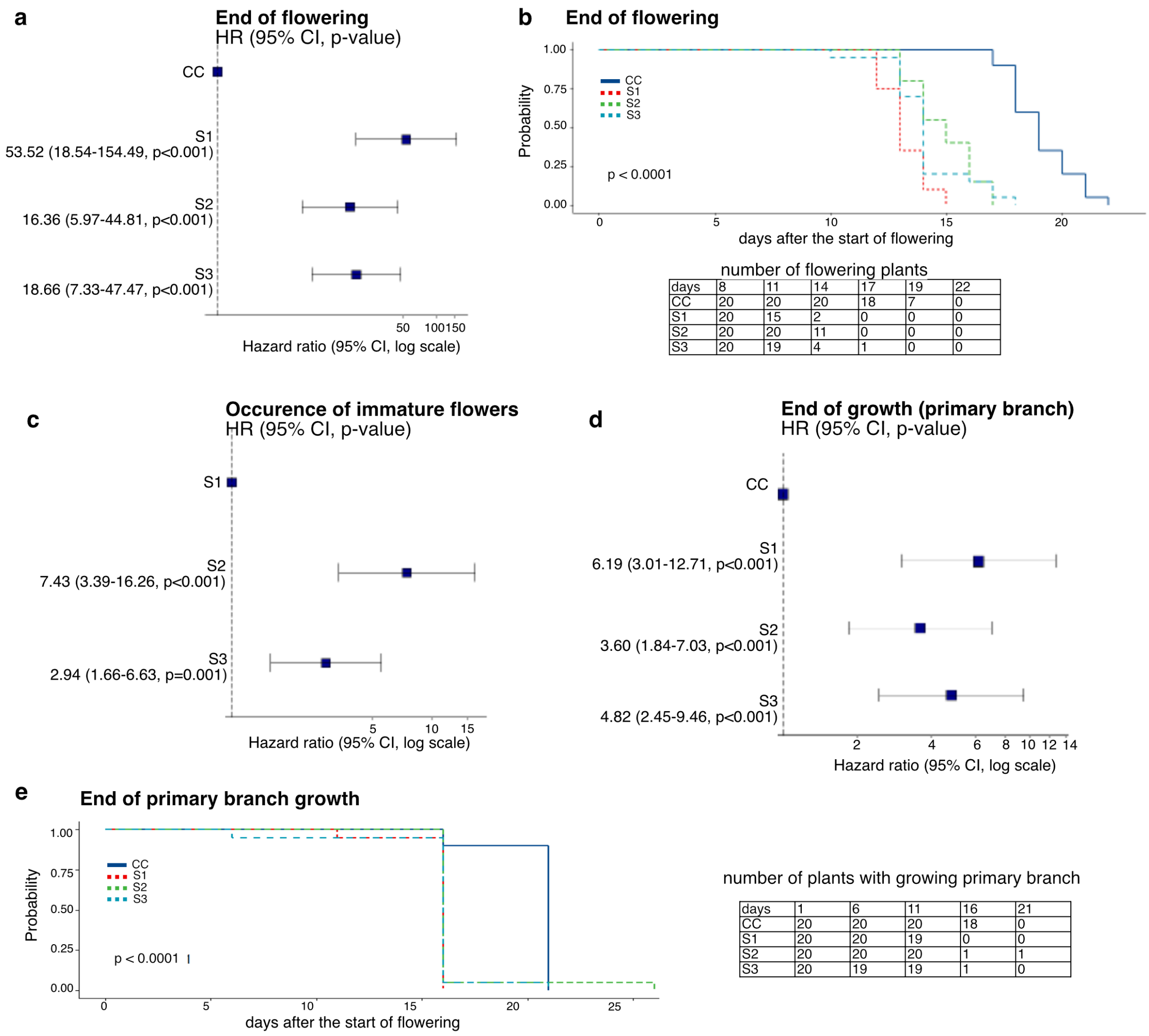


number of flowering plants

| days | 8 | 11 | 14 | 17 | 19 | 22 |
|---|---|---|---|---|---|---|
| CC | 20 | 20 | 20 | 18 | 7 | 0 |
| S1 | 20 | 15 | 2 | 0 | 0 | 0 |
| S2 | 20 | 20 | 11 | 0 | 0 | 0 |
| S3 | 20 | 19 | 4 | 1 | 0 | 0 |

number of plants with growing primary branch

| days | 1 | 6 | 11 | 16 | 21 |
|---|---|---|---|---|---|
| CC | 20 | 20 | 20 | 18 | 0 |
| S1 | 20 | 20 | 19 | 0 | 0 |
| S2 | 20 | 20 | 20 | 1 | 1 |
| S3 | 20 | 19 | 19 | 1 | 0 |

**Supplementary Figure S2.** Statistical assessment of temperature effects of flowering traits. Flowering ending (a, b), production of immature flowers (c), and end of primary branch growth (d, e) were evaluated using the Kaplan-Meier survival estimate method (survival probability), followed by Cox proportional hazard regression model to provide hazard ratio (HR) and confidence intervals (CI). When HR between CC and S1/2/3 conditions is less than 1 (HR<1), S1/2/3 is less likely to differ of CC. If HR>1 between CC and S1/2/3, S1/2/3 has a shorter survival (shorter flowering in a, and shorter growth period of the primary branch in d). (b) The probability of still flowering on given days after the start of flowering is given. The table provides the number of plants that are still flowering on given days after the start of flowering for each condition. (c) The HR indicates that S2 and S3 significantly induce the production of immature flowers. (e) The probability of primary branch growing on given days after the start of flowering is given. The table provides the number of plants with a growing primary branch on given days after the start of flowering for each condition.

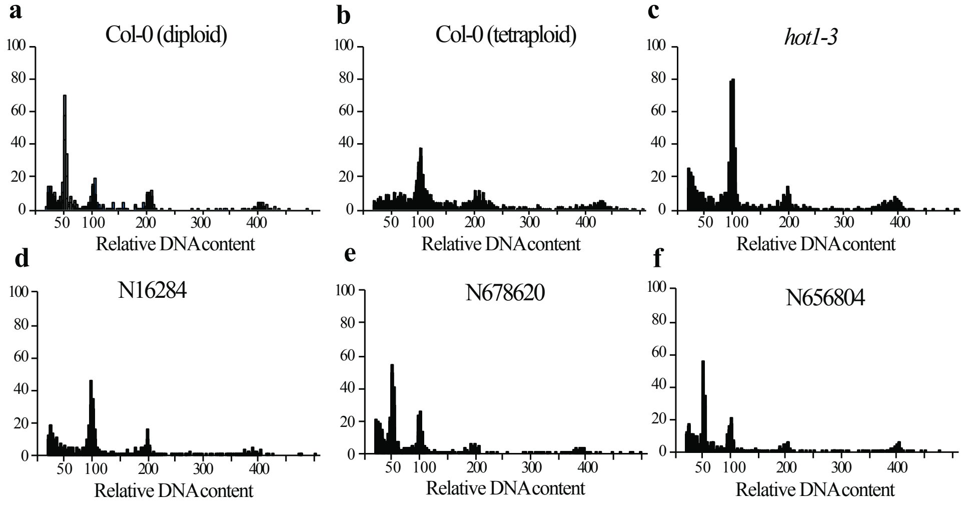


**Supplementary Figure S3.** Ploidy measurements of the *hot1* mutants.

Flow cytometry histograms showing representative profiles of peaks from diploid (a) and tetraploid (b) Col-0 and *hot1* plants grown under the CC growth condition. We compared *hot1-3* (c), the original seed stock (N16284) (d) and two different Salk lines in HSP101: N678620/SALK_066374C (e) and N656804/SALK_099583C (f). The first peak corresponds to 2C, the second to 4C and the third to 8C nuclei (8C peak not shown in tetraploid Col-0 and *hot1-3*).
