## Supplementary Tables S1-S3 for "Adaptive response to long-term high temperatures during the reproductive development in *Arabidopsis thaliana*"

**Supplementary Table S1.** Statistical comparison of the flowering production rate. Negative binomial regression estimated the flowering production rate at each condition, which was compared by ANOVA, followed by the Tukey test for multiple pairwise comparisons.

| Comparisons | estimate | S.E. | Df | t-ratio | p-value |
|---|---|---|---|---|---|
| CC vs. S1 | 0.02546 | 0.00357 | 76 | 7.134 | < 0.0001 |
| CC vs. S2 | 0.00592 | 0.00357 | 76 | 1.658 | 0.3528 |
| CC vs. S3 | 0.01944 | 0.00357 | 76 | 5.447 | < 0.0001 |
| S1 vs. S2 | -0.01955 | 0.00357 | 76 | -5.476 | < 0.0001 |
| S1 vs. S3 | -0.00602 | 0.00357 | 76 | -1.687 | 0.3375 |
| S2 vs. S3 | 0.01352 | 0.00357 | 76 | 3.789 | 0.0017 |

**Supplementary Table S2.** Statistical comparison of the primary branch growth rate. Exponential growth model estimated the growth rate for plant, which was compared by ANOVA, followed by the Tukey test for multiple pairwise comparisons.

| Comparisons | estimate | S.E. | Df | t-ratio | p-value |
|---|---|---|---|---|---|
| CC vs. S1 | 0.1764 | 0.0204 | 76 | 8.655 | < 0.0001 |
| CC vs. S2 | 0.2247 | 0.0204 | 76 | 11.024 | < 0.0001 |
| CC vs. S3 | 0.2425 | 0.0204 | 76 | 11.896 | < 0.0001 |
| S1 vs. S2 | 0.0483 | 0.0204 | 76 | 2.369 | 0.0919 |
| S1 vs. S3 | 0.0660 | 0.0204 | 76 | 3.240 | 0.0094 |
| S2 vs. S3 | 0.0178 | 0.0204 | 76 | 0.872 | 0.8195 |

**Supplementary Table S3.** List of primers used for RT-qPCR

| | |
|---|---|
| HsfA1d RT-F | GGATTCAACACCAGTGGACAATG |
| HsfA1d RT-R | AGGAGACCCATCTGTTGAGTCAG |
| HSFA2 qPCR F | TGAAGGGTTTTTAGCAGGACAA |
| HSFA2 qPCR R | CTGCAAACCCATGTTCCTCC |
| HSFA4c qPCR F NEW | CAAGCTCACTTCCACCTTTCCT |
| HSFA4c qPCR R NEW | TGTTGTTTTCGCTCCAAGCA |
| HSBP Forward | TGATTCTGCCTCCCATGTCA |
| HSBP Reverse | GCTGATATGACTGCTTTTGTCCA |
| PP2a_A3 qPCR F | AAGCGGTTGTGGAGAACATGATACG |
| PP2A_A3 qPCR R | TGGAGAGCTTGATTTGCGAAATACCG |
| TMA7 qPCR F | AGCTAAGTCAGTCTCGTCGT |
| TMA7 qPCR R | AGCAAGGAGGAAAGGCGAAA |
| APX2 qPCR F | ACTCCTTGTCAGCAAACCCGAG |
| APX2 qPCR R | CTTGATGATCCTCTCTTTCTCCCA |
| HSP101 qPCR F | ATGACCCGGTGTATGGTGCTAG |
| HSP101 qPCR R | CGCCTGCATCTATGTAAACAGTG |
| TAA1 new qpcr F | TGGCTAGGGACGAAGGAAGA |
| TAA1 new qpcr R | GCTGACTCGGACATGCTTCT |
| HSP70 qpcr F | CCGTCTTCGATGCTAAGCGTCT |

| | |
|---|---|
| HSP70 qpcr R | AACCACAATCATAGGCTTCTCACC |
