## Supplemental videos S1-S7 legend for "Adaptive response to long-term high temperatures during the reproductive development in *Arabidopsis thaliana*"

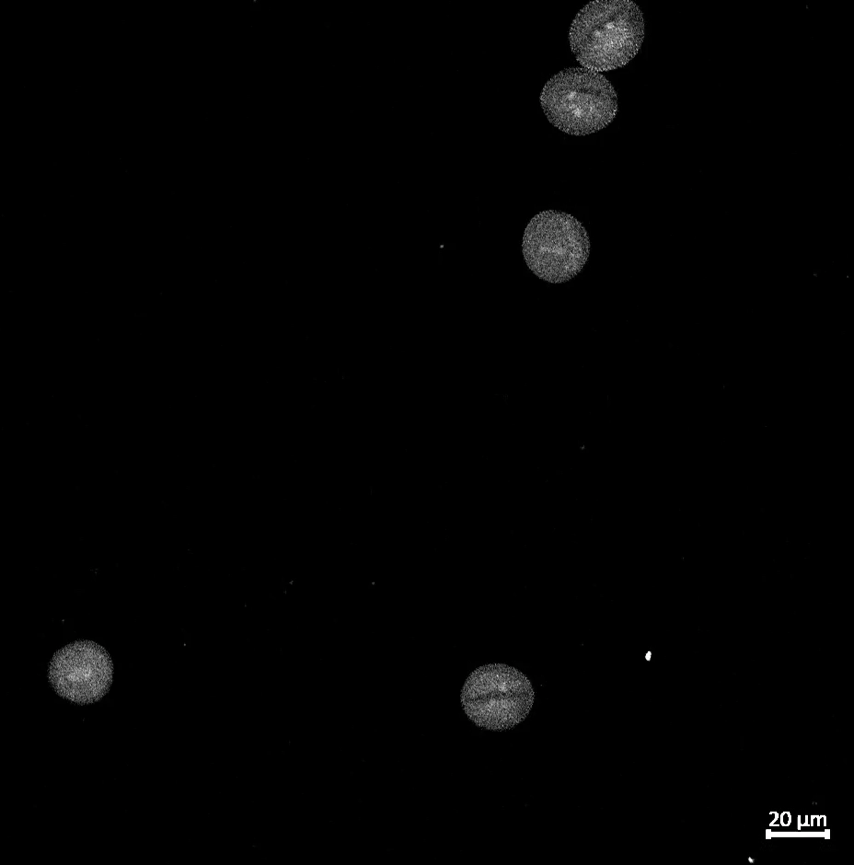

**Supplementary video S1.** Pollen grains of Col-0 plants grown under CC growth conditions. Pollen nuclei are staining by DAPI. Scale bar is 20 μm.

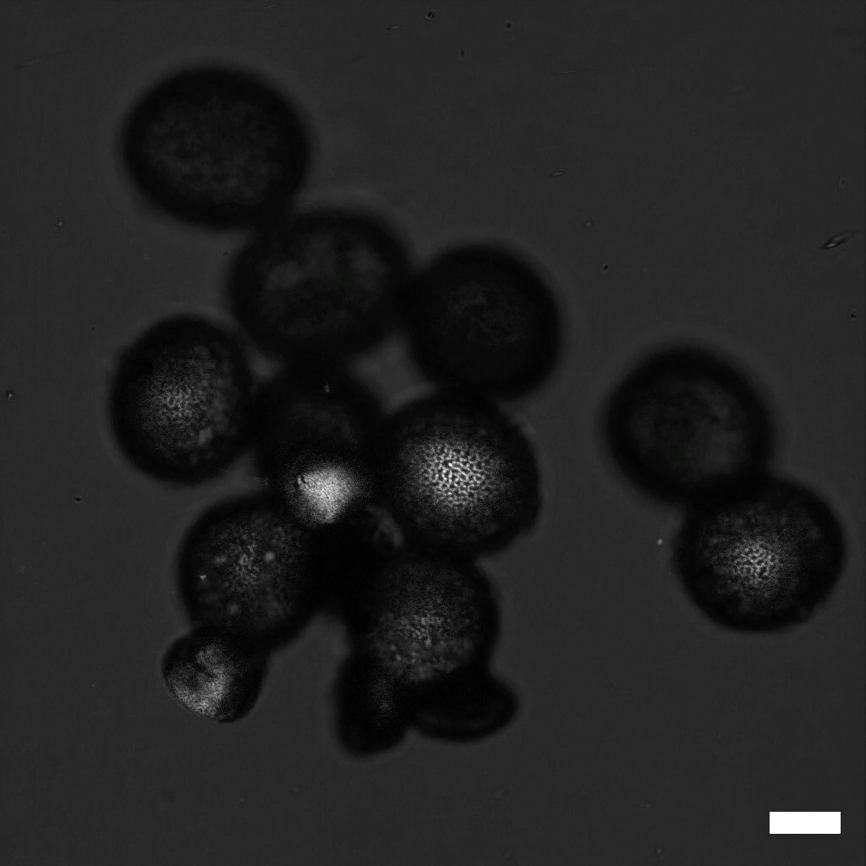

**Supplementary video S2.** Pollen grains of Col-0 plants grown under S3 growth conditions. Pollen nuclei are staining by DAPI. Scale bar is 20 μm.

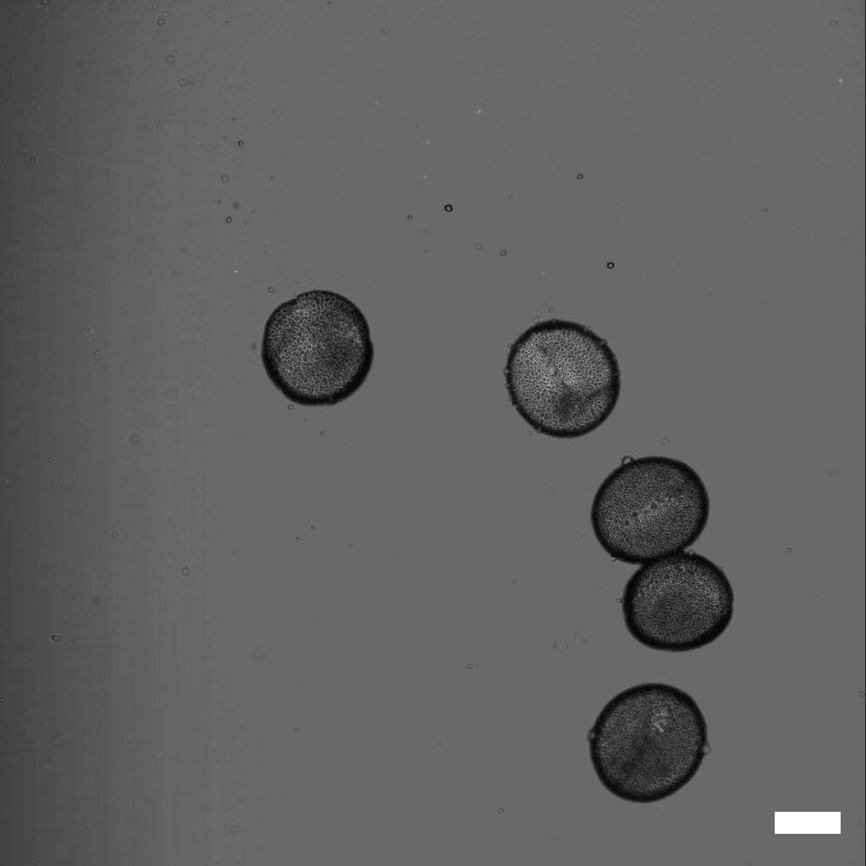

**Supplementary video S3.** Pollen grains of *hot1-3* plants grown under CC growth conditions. Pollen nuclei are staining by DAPI. Scale bar is 20 μm.

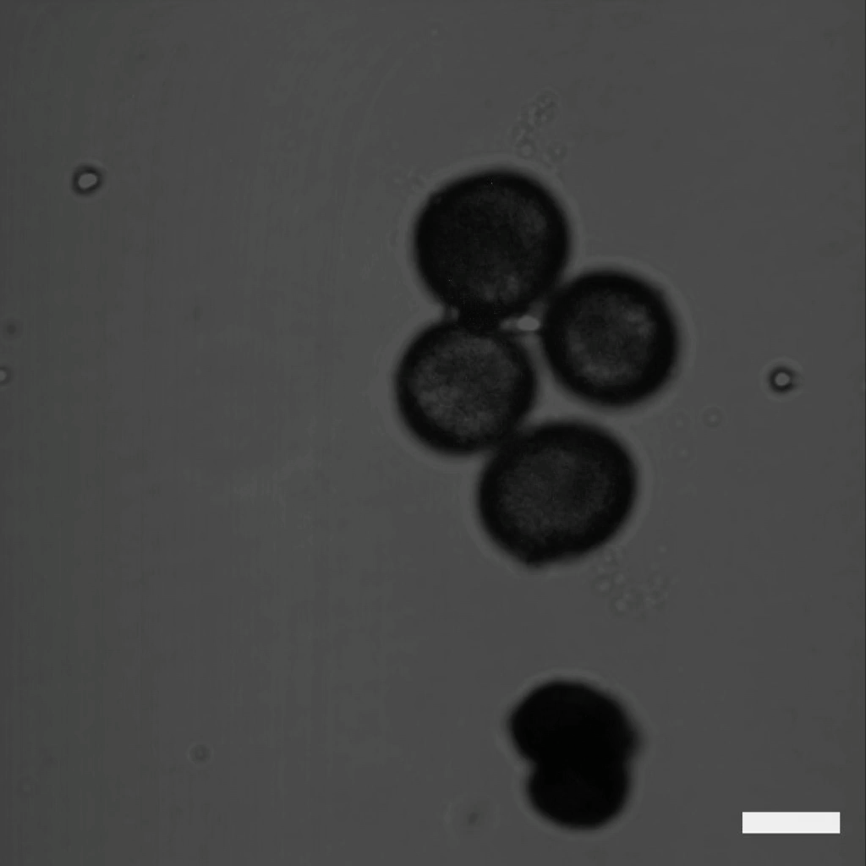

**Supplementary video S4.** Pollen grains of *hot1-3* plants grown under S3 growth conditions. Pollen nuclei are staining by DAPI. Scale bar is 20 μm.

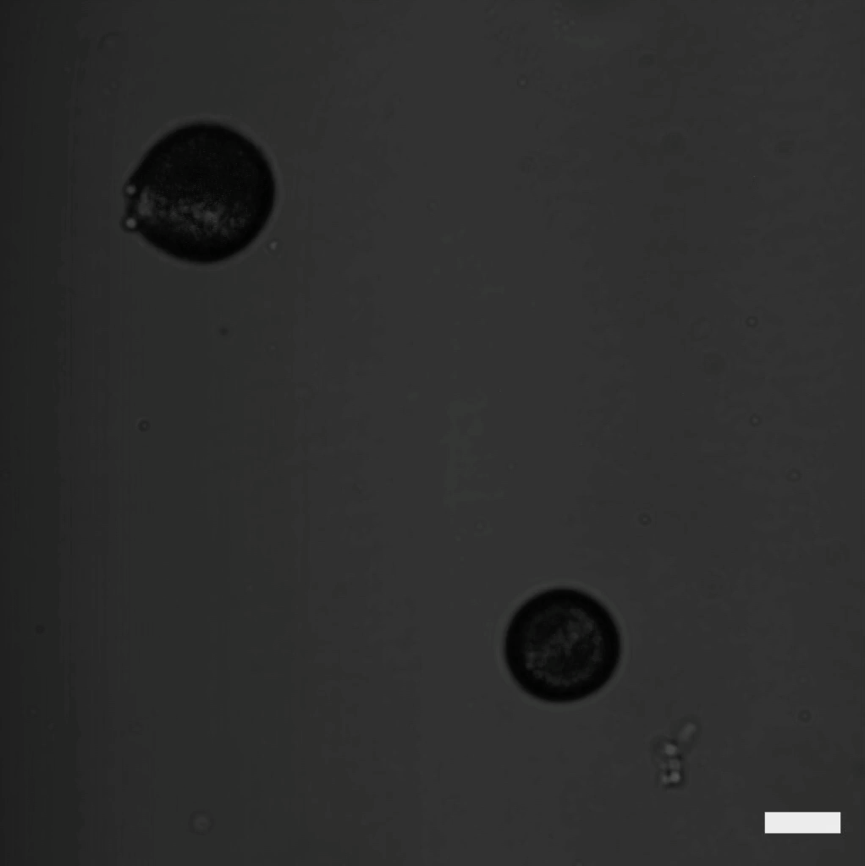

**Supplementary video S5.** Pollen grains of *hot1-3* plants grown under S3 growth conditions. Pollen nuclei are staining by DAPI. Two pollen grains of two sizes are shown. Scale bar is 20 μm.

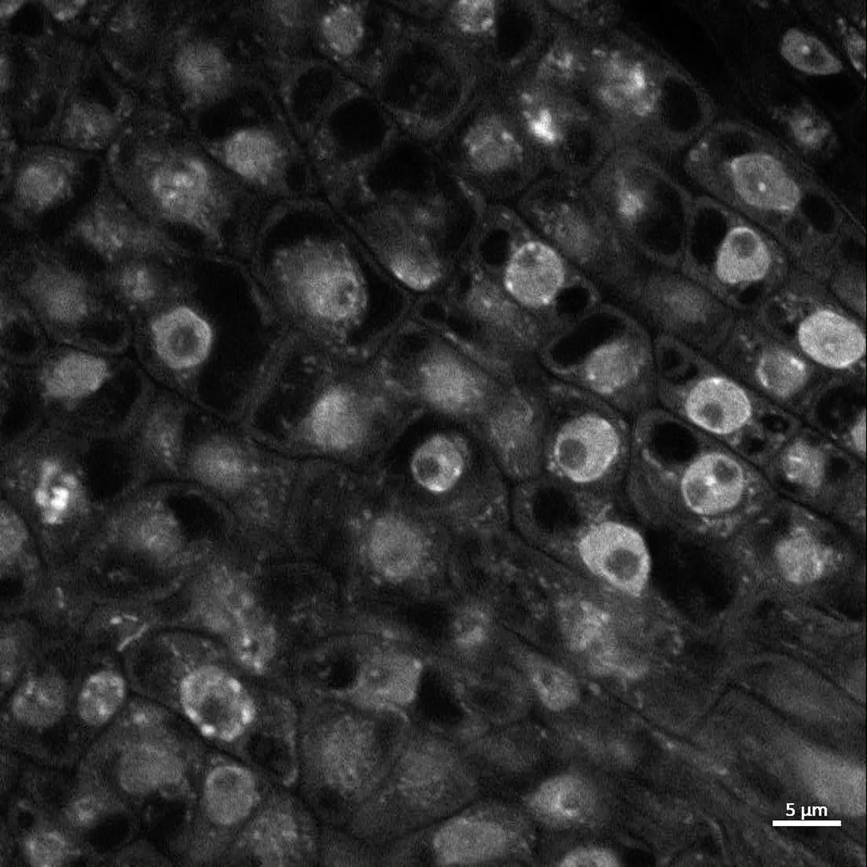

**Supplementary video S6.** Anaphase chromosomes from Col-0 sepal. Nuclei are staining by DAPI. Scale bar is 5 μm.

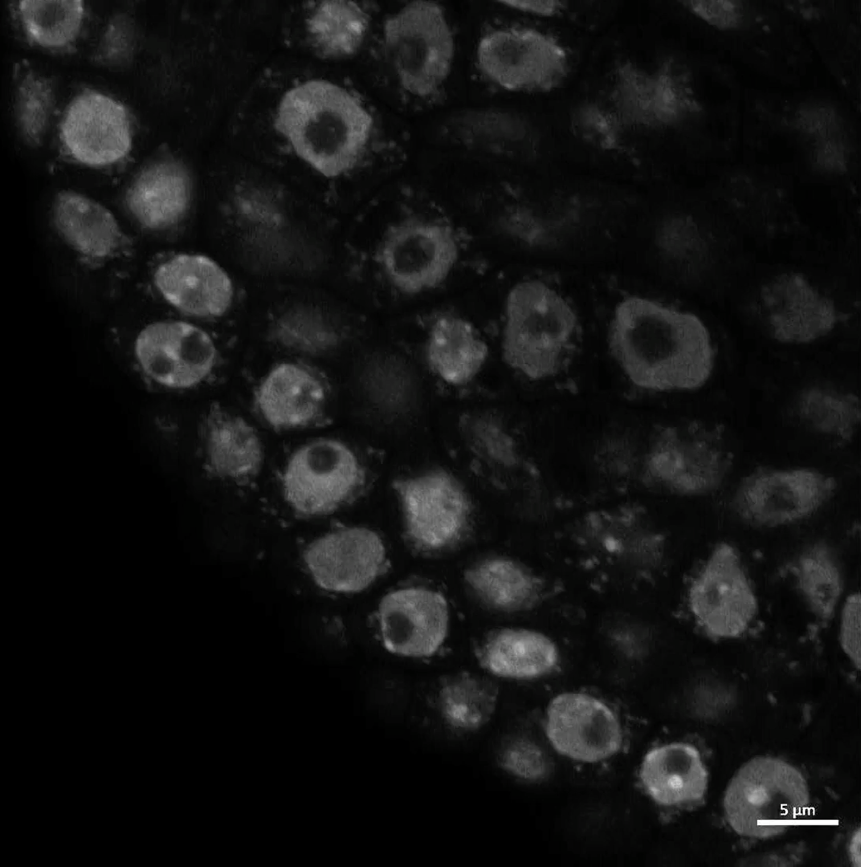

**Supplementary video S7.** Anaphase chromosomes from *hot1-3* sepal. Nuclei are staining by DAPI. Scale bar is 5 μm.
